# FER-LIKE IRON DEFICIENCY-INDUCED TRANSCRIPTION FACTOR (FIT) accumulates in homo- and heterodimeric complexes in dynamic and inducible nuclear condensates associated with speckle components

**DOI:** 10.1101/2022.12.20.520918

**Authors:** Ksenia Trofimov, Regina Gratz, Rumen Ivanov, Yvonne Stahl, Petra Bauer, Tzvetina Brumbarova

## Abstract

Some nuclear proteins undergo condensation, but the functional importance remains often unclear. The basic helix-loop-helix (bHLH) FER-LIKE IRON DEFICIENCY-INDUCED TRANSCRIPTION FACTOR (FIT) integrates internal and external signals to control iron acquisition and growth. The previously described C-terminal residues Ser271/272 allow FIT to form active complexes with subgroup Ib bHLH factors such as bHLH039. FIT has lower nuclear mobility than mutant FITmSS271AA. Here, we show that FIT undergoes a light-inducible subnuclear partitioning into nuclear condensates that we termed FIT nuclear bodies (NBs). FIT NB characteristics were examined using a standardized FIT NB analysis procedure coupled with different types of quantitative and qualitative microscopy-based approaches. FIT condensates were reversible and likely formed by liquid-liquid phase separation. FIT accumulated preferentially in FIT NBs versus nucleoplasm when engaged in protein complexes with itself and with bHLH039. FITmSS271AA, instead, localized to NBs with different dynamics. FIT colocalized with splicing and light signaling NB markers. The NB-inducing light conditions were linked with active FIT and elevated FIT target gene expression in roots. Hence, we conclude that inducible, highly dynamic FIT condensates form preferentially when transcription factor complexes are active. Inducible FIT nuclear condensates may affect nuclear mobility and integrate environmental and Fe nutrition signals.

**Highlights:**

- FIT undergoes light-induced, reversible condensation and localizes to nuclear bodies (NBs), likely via liquid-liquid phase separation
- Functionally relevant Ser271/272 defines an intrinsically disordered region and influences NB formation dynamics
- NBs are preferential sites for FIT dimerization with FIT and bHLH039, dependent on Ser271/272
- FIT NBs colocalize with NB markers related to splicing and light signaling
- Light conditions inducing NBs are linked with active FIT, in agreement with elevated FIT target gene expression in roots

## Introduction

As sessile organisms, plants adjust to an ever-changing environment and acclimate rapidly. They also control the amount of micronutrients they take up. Even though iron is one of the most abundant elements in the soil, its bioavailability as micronutrient is limited in most soils (Römheld and Marschner, 1986; Wedepohl, 1995). This poses a particular challenge for plants as they need mobilizing iron while at the same time must prevent that iron over-accumulates and becomes toxic in cells.

An essential regulatory protein needed for iron acquisition is the basic helix-loop-helix (bHLH) transcription factor (TF) FER-LIKE IRON DEFICIENCY-INDUCED TRANSCRIPTION FACTOR (FIT; Colangelo and Guerinot, 2004; Jakoby et al., 2004; Yuan et al., 2005; Bauer et al., 2007). FIT is activated upon iron deficiency downstream of a cascade of bHLH TFs (Zhang et al., 2015; Li et al., 2016b; Liang et al., 2017; Kim et al., 2019; Gao et al., 2020) and of a calcium-sensing protein kinase able to target phosphorylation site Ser271/272 of FIT (Gratz et al., 2019). FIT alone is not sufficient to upregulate iron acquisition, while it is active in a heterodimeric complex together with a member of the bHLH subgroup Ib such as bHLH039 (Yuan et al., 2008; Wang et al., 2013). Furthermore, FIT action is regulated through protein-protein contacts with multiple key players of hormonal and abiotic stress signaling pathways (Lingam et al., 2011; Le et al., 2016; Wild et al., 2016; Cui et al., 2018; Gratz et al., 2019, 2020). Thus, FIT behaves as a regulatory hub in root cells that perceives external and internal cues to adjust iron acquisition with growth (Schwarz and Bauer, 2020; Kanwar et al., 2021).

The subcellular localization of the FIT-bHLH039 module is remarkable. bHLH039 alone is inactive and present mainly close to the plasma membrane in cytoplasmic foci, while bHLH039 together with FIT localizes in the nucleus (Trofimov et al., 2019). FIT is predominately localized in the nucleus but not as mobile compared to mutant FITmSS271AA, that is a less active mutant form of FIT (Gratz et al., 2019). Subcellular partitioning of proteins involved in nutrient uptake has until now not been enough in the focus of research to understand the significance of differing subcellular localization patterns.

One prominent type of subnuclear partitioning occurs when biomolecular condensates, or nuclear bodies (NBs), form. NBs are membrane-less, nuclear sub-compartments, which can be of stable or dynamic nature. To form condensates, proteins need to have particular features that enable protein interactions and compaction in three-dimensional space. Intrinsically disordered regions (IDRs) are flexible protein regions that allow conformational changes, and thus various interactions, leading to the required multivalency of a protein for condensate formation (Tarczewska and Greb-Markiewicz, 2019; Emenecker et al., 2020). As Arabidopsis TFs are enriched in IDRs (Strader et al., 2022) it is not unlikely that the resulting multivalency in TFs drives condensation and results in microenvironments for interaction, probably more often than so far studied. IDRs are particularly characteristic in bHLH TFs in vertebrates and invertebrates (Tarczewska and Greb-Markiewicz, 2019), suggesting that this feature may also be relevant for the bHLH TFs of plants. One possibility for condensates to form is to undergo liquid-liquid phase separation (LLPS). In this process, a solution is demixed into two or more phases (Emenecker et al., 2020). This mechanism has been examined in simplified *in vitro* systems, but the involvement of different cell components renders the mechanism more complex *in vivo* (Fang et al., 2019; Riback et al., 2020; Zhu et al., 2021).

Various NBs are found. Plants and animals share several of them, e.g. the nucleolus, Cajal bodies, and speckles. The nucleolus is involved in transcription of ribosomal DNA, processing of ribosomal RNA, and ribosome biogenesis (Kalinina et al., 2018; Lafontaine et al., 2021). Nucleoli share components and function with Cajal bodies, which are e.g. ribonucleoproteins and RNA processing (Love et al., 2017; Trinkle-Mulcahy and Sleeman, 2017). Speckles are known spliceosomal sites (Reddy et al., 2012; Galganski et al., 2017). Plant-specific NBs are photobodies (PBs), which are triggered by light, temperature, and circadian clock (Pardi and Nusinow, 2021). PBs harbor regulatory complexes of the photomorphogenic responses, including photoreceptors like phytochromes (phy) and bHLH TFs belonging to the PHYTOCHROME INTERACTING FACTORs (PIFs; Pardi and Nusinow, 2021). Another trigger for inducible and reversible condensate formation is temperature (Jung et al., 2020; Zhu et al., 2021).

NBs may act as hubs integrating environmental signals (Emenecker et al., 2020; Meyer, 2020). Especially PBs may combine external cues, such as light, as an input signal to steer developmental processes (Kaiserli et al., 2015; Meyer, 2020; Pardi and Nusinow, 2021). It is proposed that the formation of NBs could be an ancient mechanism for spatial organization within the nucleus (Emenecker et al., 2020). As more evidence on condensation in plants arises, this topic remains barely examined in the scope of plant nutrition.

Here, we report a novel property of FIT, namely that FIT undergoes light-inducible and reversible nuclear condensation, which we detected as FIT nuclear bodies (NBs). The motivation for our study was then to elucidate the mechanism behind subcellular distribution and nuclear mobility of FIT and obtain functional hints. We developed a standardized FIT NB analysis procedure to characterize quantitative and qualitative aspects of the dynamic NB formation using different microscopy-based techniques. Thereby, we linked FIT NB formation with the activity status of FIT to form functional protein complexes. Splicing was also associated with the light-induced FIT NBs. Thus, this study suggests that FIT NBs are regulatory hubs steering nutritional signalling, and associating functional significance to FIT protein condensate formation.

## Results

### FIT localizes to NBs in light-inducible and dynamic manner likely as a result of LLPS

The TF FIT has a dynamic mobility and capacity to form TF complexes inside plant cells (Gratz et al., 2019; Trofimov et al., 2019). To explore possible mechanisms for dynamic FIT subcellular localization, we performed a microscopic study on FIT-GFP protein localization in the root epidermis of the root differentiation zone of 5-d-old iron-deficient seedlings of *Arabidopsis thaliana* (Arabidopsis), where FIT is active and iron acquisition occurs (FIT_pro_:FIT-GFP, this study; 35S_pro_:FIT-GFP, Gratz et al., 2019). At first microscopic inspection, FIT-GFP was evenly distributed within the nucleus (t= 0 in **Figure 1A**, FIT_pro_:FIT-GFP; t= 0 in **Supplemental Figure S1A**, 35S_pro_:FIT-GFP), as expected (Gratz et al., 2019; Trofimov et al., 2019). When whole seedlings were exposed to 488 nm laser light for several minutes, FIT became re-localized at the subnuclear level. After a lag time, FIT-GFP was present in one to four discrete spots per nucleus, visible after 40 min earliest, sometimes taking up to 120 min to appear (t= 90 minutes in **Figure 1A**, FIT_pro_:FIT-GFP; t= 40 min in **Supplemental Figure S1A**, 35S_pro_:FIT-GFP). Hence, FIT-GFP was re-localized in a blue light-inducible manner at the subnuclear level, irrespective of either of the two promoters used. During the imaging process or an additional hour of darkness, these spots disappeared (**Figure 1A**, t= 120 min), indicating that the formation of these spots was reversible. Seedlings that were kept in white light and imaged immediately did not show this localization pattern (**Supplemental Figure S1B)**. The observation of FIT nuclear spots in the root epidermis of the root differentiation zone was very interesting, suggesting that these might perhaps be NBs containing FIT. However, further inspection of the nuclear spots in root cells in this differentiating root zone was challenged by the small size and low accessibility of the nucleus, and especially considering the long lag time for detecting the nuclear spots. These factors made it impossible for us to apply quantitative fluorescence microscopy techniques to draw validated conclusions on the nature, dynamics, and functional significance of nuclear spots in root epidermis cells of the root differentiation zone of iron-deficient seedlings.

**Figure 1:**
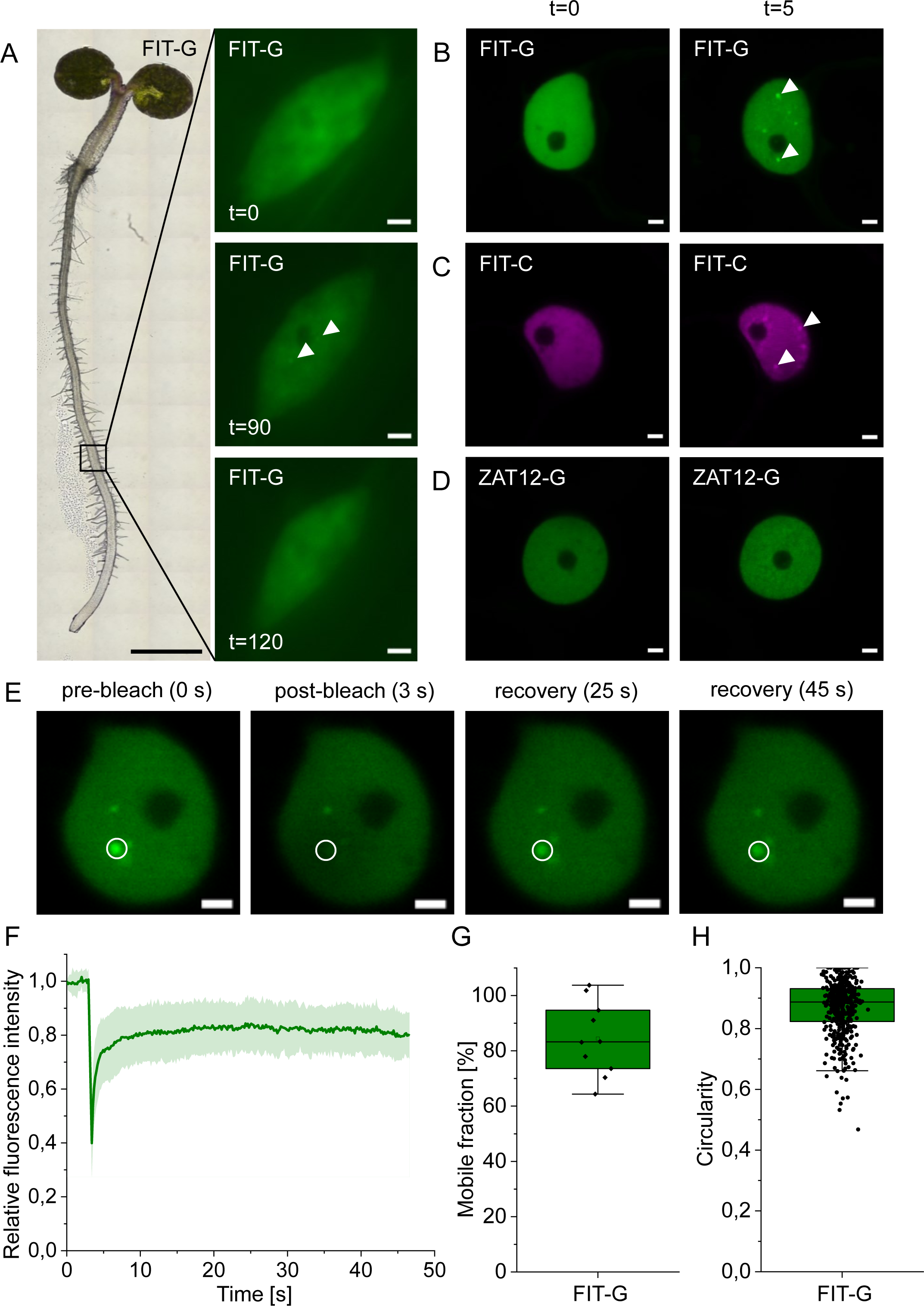
FIT accumulated in nuclear condensates, termed FIT nuclear bodies (NBs) in a light-inducible manner, most likely following liquid-liquid phase separation (LLPS). A, Induction of FIT NBs in Arabidopsis root epidermis cells of the root differentiation zone. Left, light microscopy overview image of a 5-d-old Arabidopsis seedling (proFIT:FIT-GFP) grown under iron deficiency. Right, nuclear localization of FIT-GFP in the root epidermis cells as indicated in the overview image, at t=0, t=90, and t=120 min. FIT-GFP signals were evenly distributed in the nucleus at t=0 min, and after induction by excitation with 488 nm laser NB formation accumulated in NBs at t=90 min, but also disappeared as shown at t=120 min. Note that root epidermis cells developed few NBs, sometimes taking up to two hours to appear. Three independent experiments with three plants were conducted. In the indicated region of interest, approximately 3-10 nuclei of 20 examined nuclei of the root epidermis cells showed NBs. A representative image from one nucleus is shown. B-D, Fluorescence protein analysis in transiently transformed *N. benthamiana* leaf epidermis cells. Confocal images of B, FIT-GFP, C, FIT-mCherry, and D, ZAT12-GFP at t=0 and t=5 min. At t=0 min, FIT-GFP and FIT-mCherry showed an even distribution within the nucleus. Following a 488 nm laser excitation, numerous NBs were clearly visible in all examined transformed cells at t=5 min. These NBs were termed FIT NBs. Under the same imaging conditions, ZAT12-GFP did not show NB formation. According to these results, a standardized FIT NB analysis procedure was set up (**Supplemental Figure S1C**). See also **Supplemental Movie S1A-C**. Representative images from two to three independent experiments. E-G, FRAP measurements to test for liquid-like behavior of FIT NBs, using the standardized FIT NB analysis procedure in transiently transformed *N. benthamiana* leaf epidermis cells. E, Representative images of the fluorescent signal during a FRAP experiment, taken before bleaching (0 s) and recovery of fluorescence at three time points after bleaching from 3 s to 45 s within the circled region of a NB. F, Line diagram representing the relative fluorescence during a FRAP measurement for 10 NBs, showing a high fluorescence recovery rate of FIT-GFP within NBs. Dark green line, mean value; light green filled area, variation. G, Box plot diagram representing the mobile fraction of FIT-GFP calculated based on the relative fluorescence recovery in F. The diagram indicates high mobility of FIT. The mean was calculated from 10 NBs from 10 nuclei from a transformed plant. Three independent experiments were conducted, one representative result is shown. H, Box plot diagram representing quantification of the FIT NB shape with the software ImageJ (National Institutes of Health), indicating that FIT NBs have circular shape. Mobility and circularity characteristics indicate that FIT NBs are most likely liquid condensates that are the result of LLPS. The mean was calculated from all NBs visible in 15 nuclei from a transformed plant. Two independent experiments were conducted, one representative result is shown. Box plots show 25-75 percentile with min-max whiskers, mean as small square and median as line. Scale bars of nuclei images, 2 µm; scale bar full seedling, 1 mm. Arrowheads indicate NBs. G = GFP; C = mCherry.

Since we had previously established a reliable plant cell assay for studying FIT functionality, we adapted it to study the characteristics of the prospective FIT NBs (Gratz et al., 2019, 2020; Trofimov et al., 2019). For this assay, FIT-GFP was transiently expressed in *Nicotiana benthamiana* (*N. benthamiana*) leaf epidermis cells under a β-estradiol-inducible promoter. Under such controlled protein expression condition, FIT-GFP re-localized also into nuclear spots as observed in the Arabidopsis root epidermis, again triggered by treatment with a 488 nm laser light stimulus. Differences were, however, the duration of the lag time needed to observe this phenomenon, and the number of nuclear spots. As in Arabidopsis, FIT-GFP localized initially in uniform manner to the entire nucleus (t=0) of *N. benthamiana* leaf epidermis cells (**Figure 1B**). A short duration of 1 min 488 nm laser light excitation induced the formation of FIT-GFP signals in discrete spots inside the nucleus, which became fully visible after only five minutes (t=5; **Figure 1B and Supplemental Movie S1A**). The nuclear FIT spots were systematically initiated, and nearly all nuclei in the imaged leaf disk showed numerous spots. A similar laser light excitation procedure was previously found to elicit PB formation of cryptochrome2 (CRY2) in Arabidopsis protoplasts and HEK293T cells (Wang et al., 2021). We deduced that the spots of FIT-GFP signal were indeed very likely NBs (for this reason hereafter termed FIT NBs). FIT NB formation was not dependent on the fluorescent tag, as it was similar for FIT-mCherry when co-excited with 488 nm laser light (**Figure 1C**). Another TF and interactor of FIT, ZINC FINGER OF ARABIDOPSIS THALIANA12-GFP (ZAT12-GFP; Le et al., 2016), did not localize to NBs under the same imaging conditions (**Figure 1D and Supplemental Movie S1B**). Therefore, we concluded that FIT localization to NBs was a specificity of FIT and that formation of FIT NBs was not artificially caused by fluorescent tags or the imaging setup. Importantly, the *N. benthamiana* epidermis expression system was suited to control the parameters for light-induced triggering of FIT NBs and their quantitative analysis by fluorescence microscopy. We then developed a standardized experimental procedure for qualitative and quantitative FIT NB analysis in *N. benthamiana* (hereafter named ‘standardized FIT NB analysis procedure’; **Supplemental Figure S1C**).

Liquid-liquid phase separation (LLPS) is a possible way for condensate formation, and liquid-like features are quantifiable by mobility and shape analysis within condensates (Shin et al., 2017; Wang et al., 2021). We used the standardized FIT NB analysis procedure to examine whether this could also be a mechanism underlying the FIT NB formation. Mobility of FIT NBs was tested with the fluorescence recovery after photobleaching (FRAP) approach (Bancaud et al., 2010; Trofimov et al., 2019) by recording the recovery of the fluorescence intensity over time in a bleached NB (**Figure 1E-G**). According to relative fluorescence intensity the fluorescence signal recovered rapidly within FIT NBs (**Figure 1F**), and the calculated mobile fraction of the NB protein was on average 80% (**Figure 1G**). Shape analysis of FIT NBs showed that the NBs reached a high circularity score (**Figure 1H**). According to Wang et al. (2021), fluorescence recovery and circularity scores as the ones measured for FIT NBs reflect high mobility and circular shape. Thus, FIT NBs behave in a liquid-like manner suggesting that LLPS might be the mechanism leading to FIT NB formation.

In summary, FIT-GFP forms light-inducible FIT NBs in roots and upon transient expression in *N. bethamiana* leaf epidermis cells. The developed standardized FIT NB analysis procedure was well suited for investigating dynamic properties of light-induced FIT NBs and characterizing them as the likely result of LLPS. Because of these properties, it is justified to term them ‘FIT NBs’. We hypothesized that NB formation is a feature of the FIT protein that provides regulatory specificity, and we subsequently investigated this hypothesis using the developed standardized FIT NB analysis procedure in all subsequent assays below.

### FIT forms homodimeric complexes preferentially in NBs, dependent on Ser271/272

Next, we asked which properties of the FIT protein enable NB formation. Residue Ser271/272 is important for the homo- and heterodimerization capacity of FIT (Gratz et al., 2019). We therefore asked whether this site has an influence on FIT NB formation, and we compared the ability for NB formation of phospho-mutated FIT with that of wild-type FIT-GFP protein. In the work of Gratz et al. (2019), the phospho-mimicking FITmS272E protein did not show significant changes in its behavior compared to wild-type FIT. However, the FITmSS271AA-GFP mutant was less able to mediate induction of iron deficiency responses and its properties differed from wild-type FIT-GFP (Gratz et al., 2019). Hence, we focused on comparing the nuclear NB-forming characteristics between wild-type FIT-GFP and FITmSS271AA-GFP mutant.

FITmSS271AA-GFP also localized to NBs. The formation of FITmSS271AA NBs was delayed in time (**Figure 2A**; t=15 instead of t=5). While FIT-GFP NB formation started in the first minutes after excitation and was fully present after 5 min (**Supplemental Movie S1A**), FITmSS271AA-GFP NB formation occurred earliest 10 min after excitation and was fully visible after 15 min (**Supplemental Movie S1C**). In addition, NB number and size of FITmSS271AA-GFP were decreased in comparison to the ones from wild-type FIT-GFP (**Figure 2, B and C**). Hence, the dynamics of NB formation were dependent on Ser271/272.

**Figure 2:**
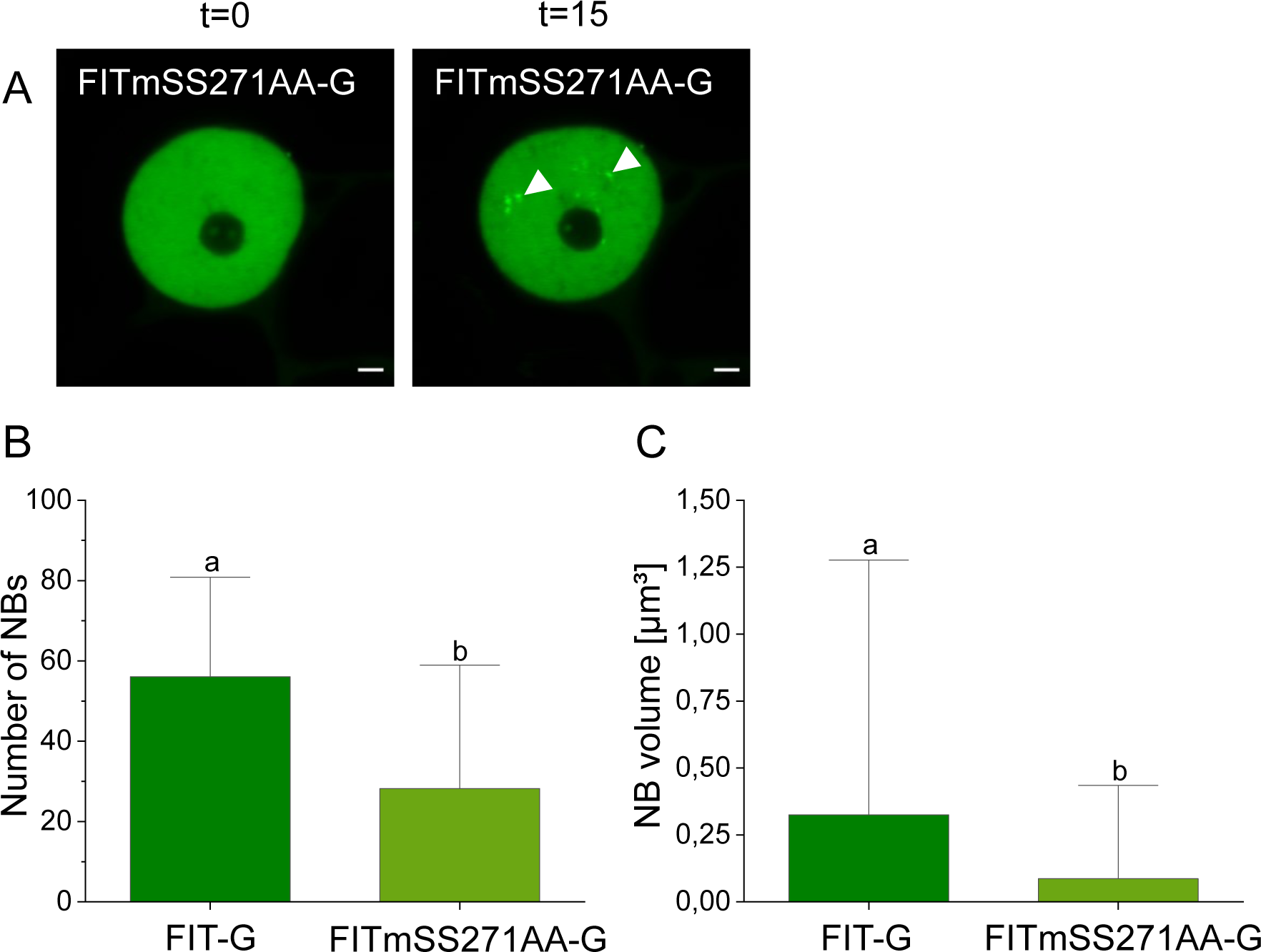
The FIT C-terminal Ser271/272 site was important for the capacity of FIT to localize to NBs. A, Confocal images of nuclear localization of FITmSS271AA-GFP at t=0 and t=15 min. FITmSS271AA-GFP accumulated in NBs, but NB formation required a longer time compared to FIT-GFP. See also Supplemental Movie S1, A and C. Two independent experiments. Representative images from one nucleus. B, Bar diagrams showing in B, number of NBs, and in C, the sizes of NBs with FIT-GFP and FITmSS271AA-GFP at t=5/15 min. NB number and size were determined with the software ImageJ (National Institute of Health). FIT-GFP accumulated in more and larger NBs than FITmSS271AA-GFP. See Supplemental Movie S1, A and C. FITmSS271AA-GFP lacks IDR^Ser271/272^. This IDR may be relevant for FIT NB formation (Supplemental Figure S2). In B and C, bar diagrams represent the mean and standard deviation for a quantification of 15 nuclei from a transformed plant (n = 15). Two experiments were conducted, one representative result is shown. Statistical analysis was performed with the Mann-Whitney test. Different letters indicate statistically significant differences (P < 0.05). Scale bar: 2 µm. Arrowheads indicate NBs. G = GFP. Analysis was conducted in transiently transformed *N. benthamiana* leaf epidermis cells, following the standardized FIT NB analysis procedure.

The process of condensation is facilitated when proteins possess IDRs, since, importantly, IDRs may engage in numerous interactions in space due to rapid conformational changes (Tarczewska and Greb-Markiewicz, 2019; Emenecker et al., 2020). The three-dimensional conformation of wild-type FIT had predicted stretches of intrinsic disorganization, peaking before and at the basic region of the bHLH domain, and two in the C-terminal part, one of them around the Ser271/272 site (termed IDR^Ser271/272^; **Supplemental Figure S2A**). In contrast, in the FITmSS271AA mutant this C-terminal region was no longer classified as IDR (**Supplemental Figure S2B**). This underlined the significance of the Ser271/272 site, not only for interaction (Gratz et al., 2019) but also for FIT NB formation (**Figure 2**).

We then tested whether FIT homodimerization was preferentially associated with NB formation. For that, we investigated whether FIT-GFP shows a differentiated homodimerization strength, first, inside the NBs versus the nucleoplasm (NP), and second, as wild-type FIT versus the mutant FITmSS271AA-GFP protein by performing anisotropy (or homo-FRET) measurements. Energy transfer between the same kind of fluorescently tagged proteins leads to depolarization of the emitted light (Stahl et al., 2013; Weidtkamp-Peters et al., 2022). Fluorescence anisotropy (FA) describes this depolarization and gives hints on the dimerization and oligomerization status of a protein as the FA value decreases (**Figure 3A**). We measured FA before (t=0) and after NB formation (t=5 for FIT and t=15 for FITmSS271AA), and analyzed the homodimerization strength for the whole nucleus, the NBs, and the residual NP (**Figure 3B-D**). Free GFP and GFP-GFP constructs were used as references for monomers and dimers (**Figure 3C and D**).

**Figure 3:**
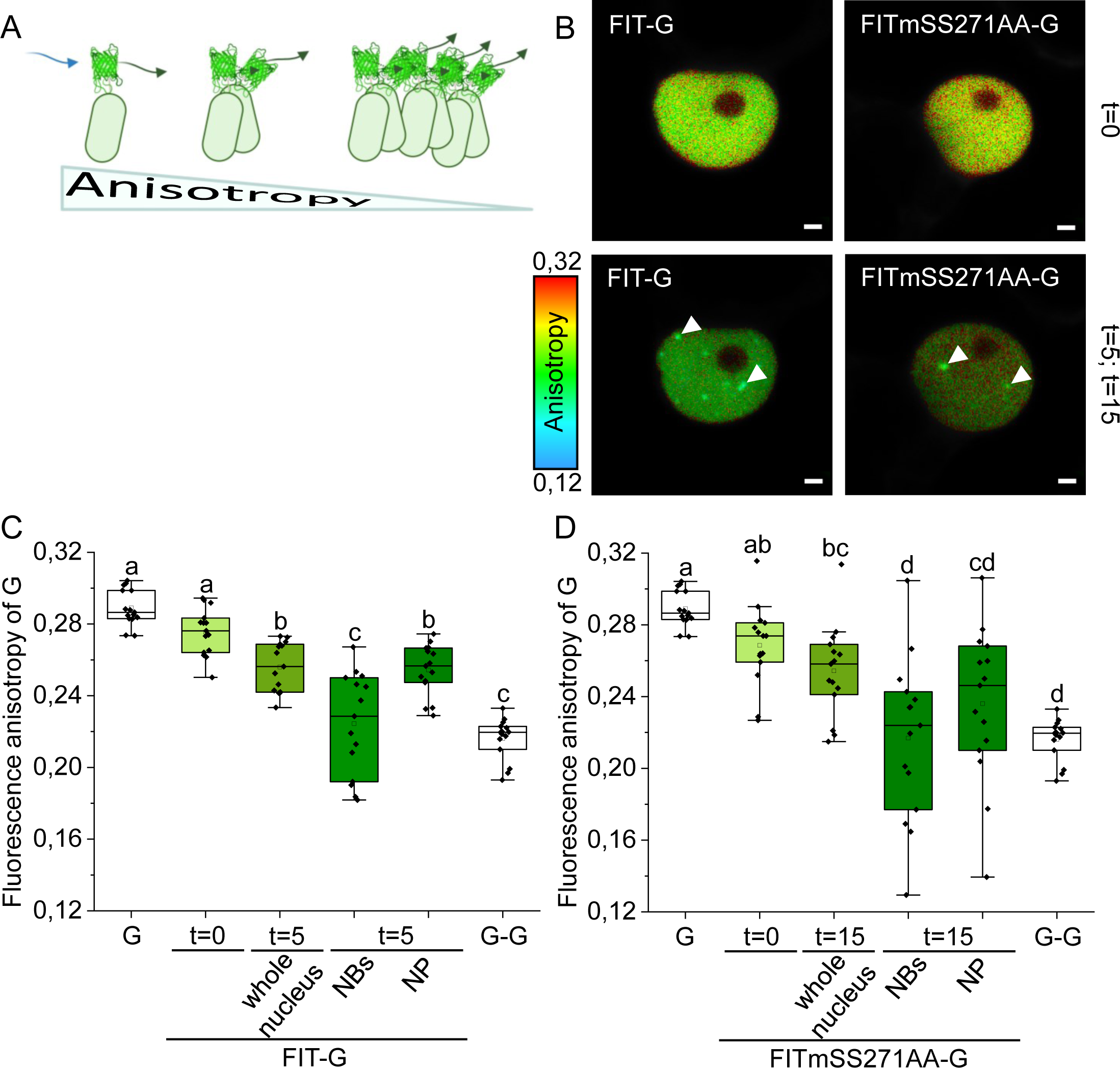
FIT was present in homodimeric protein complexes in NBs, dependent on Ser271/272 site. Anisotropy (or homo-FRET) measurements of FIT-GFP and FITmSS271AA-GFP to determine homodimerization strength. A, Schematic illustration of the anisotropy principle. Energy transfer between the same kind of fluorescently tagged proteins leads to depolarization of the emitted light. Extent of the depolarization gives a hint on dimerization and oligomerization of a protein as the fluorescence anisotropy (FA) value decreases. B, Representative images showing color-coded FA values of FIT-GFP and FITmSS271AA at t=0 and t=5/15 min. C-D, Box plots representing quantification of FA values. FA was measured at t=0 within the whole nucleus and at t =5/15 min within the whole nucleus, in NBs and in residual NP. Free GFP and GFP-GFP served as references for mono- and dimerization. FA values for C, FIT-GFP, and D, FITmSS271AA-GFP. In C and D, FA values were calculated from 10-15 nuclei from a transformed plant (n = 10-15). Two experiments were conducted, one representative result is shown. C and D show the same free GFP and GFP-GFP references because both measurements were performed on the same day. FA values decreased for FIT-GFP, but not FITmSS271AA-GFP, in the whole nucleus (compare t=0 with t=5/15 min). FA values were also lowered in NBs versus NP in the case of FIT-GFP but not FITmSS271AA-GFP (compare t=5/15 min of NBs and NP). This indicates stronger homodimerization of FIT than FITmSS271AA-GFP in the whole nucleus and in NBs. IDR^Ser271/272^ may therefore be relevant for FIT NB formation and FIT homodimerization (**Supplemental Figure S2**). Box plots show 25-75 percentile with min-max whiskers, mean as small square and median as line. Statistical analysis was performed with one-way ANOVA and Tukey post-hoc test. Different letters indicate statistically significant differences (P < 0.05). Scale bar: 2 µm. Arrowheads indicate NBs. G = GFP. Fluorescence protein analysis was conducted in transiently transformed *N. benthamiana* leaf epidermis cells, following the standardized FIT NB analysis procedure.

Whole nucleus FA values were lower at t=5 than at t=0 for FIT-GFP. Additionally, FA values were lower within the NBs compared to the NP (**Figure 3C**). Compared to wild-type FIT-GFP, FA values were not reduced for mutant FITmSS271AA-GFP at t=15 compared to t=0. Also, the FA values did not differ between NBs and NP for the mutant protein and did not show a clear separation in homodimerizing/non-dimerizing regions (**Figure 3D**) as seen for FIT-GFP (**Figure 3C**). Both NB and NP regions showed that homodimers occurred very variably in FITmSS271AA-GFP.

In summary, wild-type FIT could be partitioned properly between NBs and NP compared to FITmSS271AA mutant and rather form homodimers, presumably due its IDR^Ser271/272^ at the C-terminus. NBs were nuclear sites in which FIT formed preferentially homodimeric protein complexes.

### FIT-bHLH039 interaction complexes preferentially accumulate in FIT NBs

FIT engages in protein-protein interactions with bHLH039 to steer iron uptake target gene induction in the nucleus, while mutant FITmSS271AA protein is less active in interacting with bHLH039 (Gratz et al., 2019). Hence, we tested whether FIT also interacts with bHLH039 preferentially inside NBs and whether mutant FITmSS271AA differs in this ability from wild-type FIT protein. bHLH039 alone does not localize inside the nucleus but requires FIT for nuclear localization (Trofimov et al., 2019), so that bHLH039 was not used alone to test its subnuclear localization.

Upon co-expression, FIT-GFP and bHLH039-mCherry colocalized fully in NBs that resembled the previously described FIT NBs. In the beginning, both proteins were uniformly distributed within the nucleus (t=0), and later became localized in NBs (t=5; **Figure 4A**).

**Figure 4:**
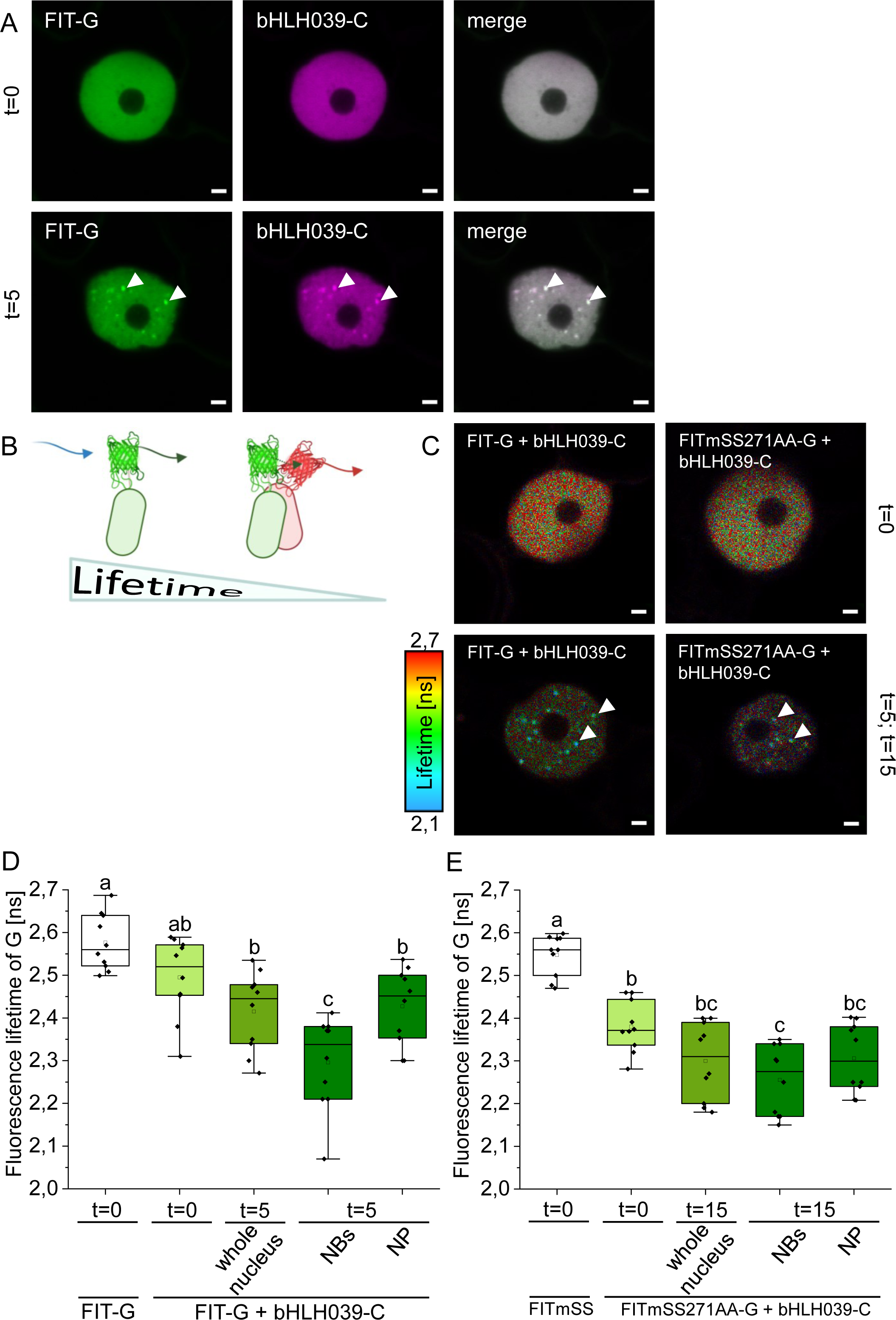
FIT was present in heterodimeric protein complexes with bHLH039 in NBs, dependent on Ser271/272 site. A, Confocal images with colocalization of FIT-GFP and bHLH039-mCherry in the nucleus. Both proteins were evenly distributed within the nucleus at t=0 and colocalized fully in FIT NBs at t=5 min. Two independent experiments with two plants each. In all examined cells, the proteins showed full colocalization. Representative images from one nucleus. B-E, FRET-FLIM measurements to determine heterodimerization strength of FIT and FITmSS271AA with bHLH039, respectively. FIT-GFP and FITmSS271AA-GFP (donor only) served as negative controls. B, Schematic illustration of the FRET-FLIM principle. Energy transfer occurs between two different fluorophores. One fluorophore acts as the donor and the other as the acceptor of the energy. In case of interaction (close proximity, ≤10 nm) the fluorescence lifetime of the donor decreases. C, Representative images showing color-coded fluorescence lifetime values of FIT-GFP and FITmSS271AA-GFP co-expressed with bHLH039-mCherry at t=0 and t=5/15 min. D-E, Box plots diagrams representing FRET-FLIM measurements at t=0 within the whole nucleus and at t=5/15 min within the whole nucleus, inside NBs and in residual NP. Lifetime values represent measurements of 10 nuclei from a transformed plant (n = 10). Two experiments were conducted, one representative result is shown. Fluorescence lifetime was reduced for the pair of FIT-GFP and bHLH039-mCherry in NBs versus NP at t=5 min, indicating protein interaction preferentially inside NBs. Fluorescence lifetime values were not significantly different for the pair FITmSS271AA-GFP and bHLH039-mCherry in this same comparison at t=15 min, indicating that this pair did not preferentially interact in NBs. IDR^Ser271/272^ may therefore be relevant for FIT NB formation, and FIT homo- and heterodimerization (**Supplemental Figure S2**). Box plots show 25-75 percentile with min-max whiskers, mean as small square and median as line. Statistical analysis was performed with one-way ANOVA and Tukey post-hoc test. Different letters indicate statistically significant differences (P < 0.05). Scale bar: 2 µm. Arrowheads indicate NBs. G = GFP; C = mCherry. Fluorescence protein analysis was conducted in transiently transformed *N. benthamiana* leaf epidermis cells, following the standardized FIT NB analysis procedure.

We then examined the heterodimerization strength of FIT-GFP and bHLH039-mCherry, and FITmSS271AA-GFP and bHLH039-mCherry by FRET-fluorescence lifetime imaging microscopy (FRET-FLIM) measurements. In case of protein interaction (close proximity, ≤10 nm), energy transfer between a fluorescently tagged donor and a fluorescently tagged acceptor decreases the fluorescence lifetime of the donor (**Figure 4B**; Borst and Visser, 2010; Weidtkamp-Peters and Stahl, 2017). We quantified the fluorescence lifetime of FIT-GFP and FITmSS271AA-GFP respective of heterodimerization before (t=0) and after NB formation (t=5 for FIT and t=15 for FITmSS271AA) in the whole nucleus, in NBs, and in the NP (**Figure 4C-E**). FIT-GFP and FITmSS271AA-GFP (donor only) served as negative controls.

Fluorescence lifetime was decreased for the pair FIT-GFP and bHLH039-mCherry at t=5 within NBs compared to all other measured areas (**Figure 4D**). In contrast to that, the fluorescence lifetime decreased for the pair FITmSS271AA-GFP and bHLH039-mCherry at t=15 was not different between NBs and NP (**Figure 4E**). This indicated that heterodimeric complexes accumulated preferentially in FIT NBs.

In summary, heterodimerization of FIT with bHLH039 was spatially concentrated in NBs versus the remaining nuclear space and was less prominent for FITmSS271AA. Hence, the capacity of FIT to form an active TF complex was coupled with its presence in NBs. The occurrence of FIT homo- and heterodimerization preferentially in NBs suggests that FIT protein interaction may drive condensation. We therefore concluded that FIT NBs may be sites with active TF complexes for iron deficiency response regulation.

### FIT NBs colocalize with speckle components

Numerous NB types are known, and they are associated with particular proteins that are indicative of the NB type. To further understand the identity, dynamics, and function of FIT NBs, we co-expressed FIT-GFP with seven different NB markers from The Plant Nuclear Marker collection (NASC) and observed NB formation and protein colocalization before (t=0) and after FIT NB formation (t=5). In cases where we detected a colocalization with FIT-GFP, we analyzed the localization of NB markers also in the single expression at t=0 and at t=5 after the 488 nm excitation, to detect potentially different patterns in single and co-expression.

All seven NB markers were expressed together with FIT-GFP, and according to the resulting extent of colocalization we subdivided them into three different types. The first type (type I) did not colocalize with FIT-GFP neither at t=0 nor at t=5 (**Supplemental Figure S3**). This was the case for the Cajal body markers coilin-mRFP and U2 snRNP-specific protein U2B”-mRFP (**Supplemental Figure S3**; Lorković et al., 2004; Collier et al., 2006). Coilin-mRFP localized into a NB within and around the nucleolus (**Supplemental Figure S3A**). The NBs of U2B”-mRFP were also close to the nucleolus (**Supplemental Figure S3B**). Hence, FIT-GFP was not associated with Cajal bodies.

The second type (type II) of NB markers were partially colocalized with FIT-GFP. This included the speckle components ARGININE/SERINE-RICH45-mRFP (SR45) and the serine/arginine-rich matrix protein SRm102-mRFP (**Figure 5**). SR45 is involved in splicing and alternative splicing and is part of the spliceosome in speckles (Ali et al., 2003), and was recently found to be involved in splicing of iron homeostasis genes (Fanara et al., 2022). SRm102 is a speckle component (Kim et al., 2016). SR45-mRFP localized barely in the NP but inside few and very large NBs that remained constant at t=0 and t=5. FIT-GFP did not colocalize in those NBs at t=0, however, it colocalized with the large SR45-mRFP NBs at t=5 (**Figure 5A**). FIT-GFP also localized in typical FIT NBs in the residual NP at t=5 (**Figure 5A**). SRm102-mRFP showed low expression in the NP and stronger expression in a few NBs that also remained constant at t=0 and t=5. FIT-GFP colocalized with SRm102-mRFP in only few instances at t=5, but not t=0, while most FIT NBs did not colocalize with SRm102-mRFP NBs (**Figure 5B**). Both SR45-mRFP and SRm102-mRFP had the same localization pattern at t=0 and t=5, irrespective of FIT-GFP co-expression or 488 nm excitation (**Supplemental Figure S4**). These type II NB markers seemed to recruit FIT-GFP into NBs after 488 nm excitation that were present (pre-existing) before FIT-GFP NB formation, while FIT-GFP localized additionally in separate FIT NBs. Hence, FIT became associated with splicing components and speckles upon the light trigger.

**Figure 5:**
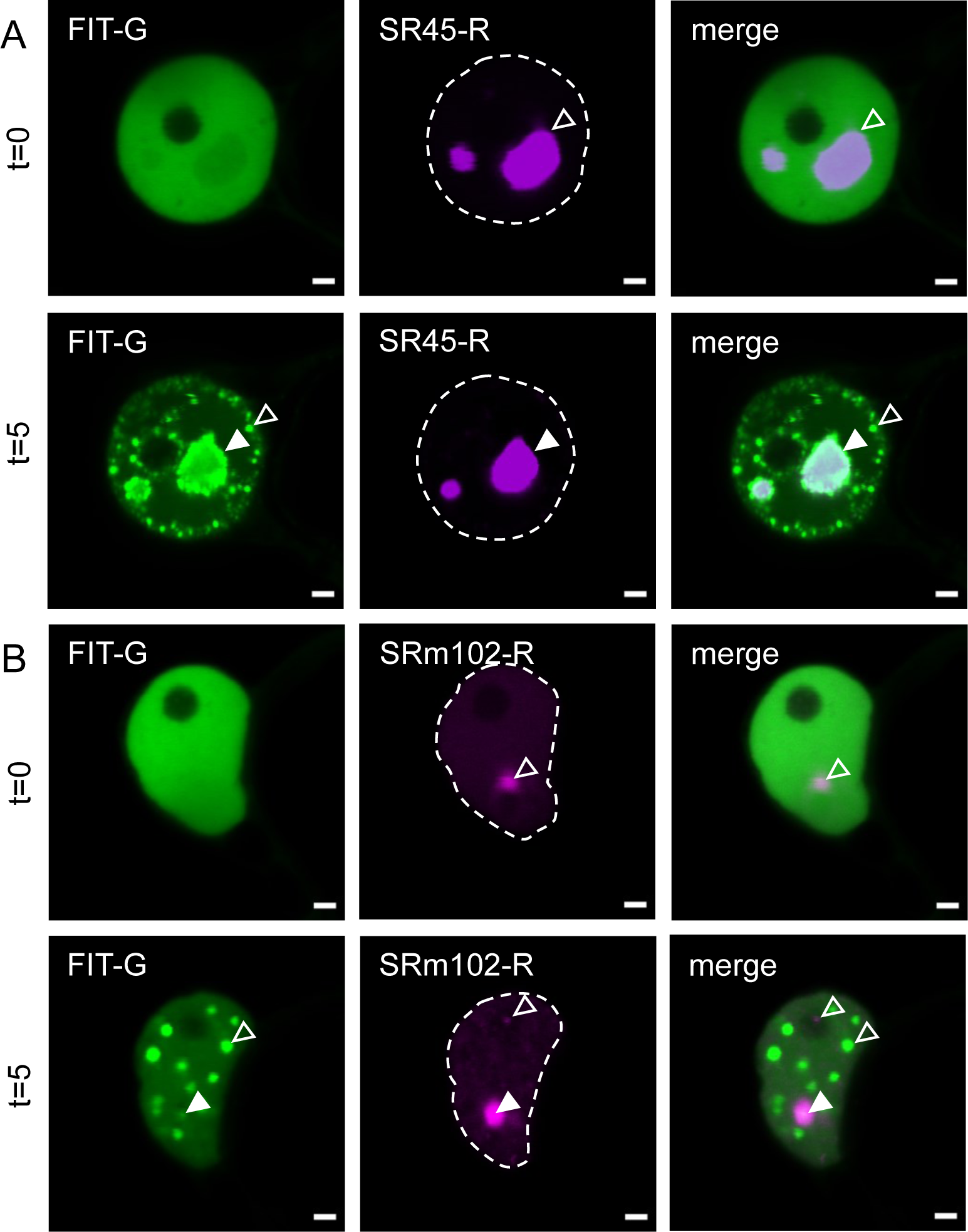
Two NB markers and splicing components were present in NBs (designated type II), revealing the dynamics of FIT to accumulate in NBs. Confocal images showing localization of FIT-GFP and NB markers (type II) upon co-expression in the nucleus at t=0 and t=5 min. Co-expression of FIT-GFP with A, SR45-mRFP, and B, SRm102-mRFP. Type II NB markers localized inside NBs at t=0 and t=5 min. Similar localization patterns were observed upon single expression, showing that SR45 and SRm102 are present in distinct NB types (compare with **Supplemental Figure S4, A and B**). FIT-GFP colocalized with type II markers in their distinct NBs at t=5 min, but not t=0. FIT-GFP additionally localized in FIT NBs at t=5 min. Type II markers were not present in FIT NBs, while FIT-GFP became recruited into the distinct type II NBs upon the light trigger. Hence, FIT NBs could be associated with speckle components. Scale bar: 2 µm. Filled arrowheads indicate colocalization in NBs, empty arrowheads indicate no colocalization in NBs. G = GFP; R = mRFP. Fluorescence protein analysis was conducted in transiently transformed *N. benthamiana* leaf epidermis cells, following the standardized FIT NB analysis procedure. In all examined cells, the proteins showed partial colocalization. Representative images from two to five independent experiments are shown. For data with type I markers (no colocalization) and type III markers (full colocalization) see **Supplemental Figure S3** and **Figure 6**.

A third type (type III) of three NBs markers, namely UAP56H2-mRFP, P15H1-mRFP, and PININ-mRFP, were fully colocalized with FIT-GFP (**Figure 6**). Until now, these NB marker proteins are not well described in plants. UAP56H2 is a RNA helicase, which is involved in mRNA export (Kammel et al., 2013). P15H1 was found as a putative Arabidopsis orthologue of an exon junction complex component in humans (Pendle et al., 2005), while PININ has a redundant role to its paralogue apoptotic chromatin condensation inducer in the nucleus (ACINUS) in alternative splicing (Bi et al., 2021). UAP56H2-mRFP and P15H1-mRFP did not localize in NBs and were not responsive to the 488 nm excitation when expressed alone or together with FIT-GFP at t=0 (**Figure 6, A and B and Supplemental Figure S4, C and D**). When co-expressed with FIT-GFP and following the 488 nm excitation, at t=5, the two NB markers adopted the FIT NB pattern and colocalized with FIT-GFP in FIT NBs (**Figure 6, A and B**). PININ-mRFP was also uniformly distributed in the nucleus at t=0 like FIT-GFP and fully colocalized with FIT NBs at t=5 (**Figure 6C**). But curiously, PININ-mRFP showed a very different localization in the single expression. Predominately, it localized to a very large NB besides several small NBs with no expression in the NP at t=0 and at t=5 (**Supplemental Figure S4E**). Thus, FIT-GFP recruited these type III NB marker and speckle proteins fully into FIT NBs. Since type III NB markers are also potentially involved in splicing and mRNA export from the nucleus, these same functions may be relevant in FIT NBs.

**Figure 6:**
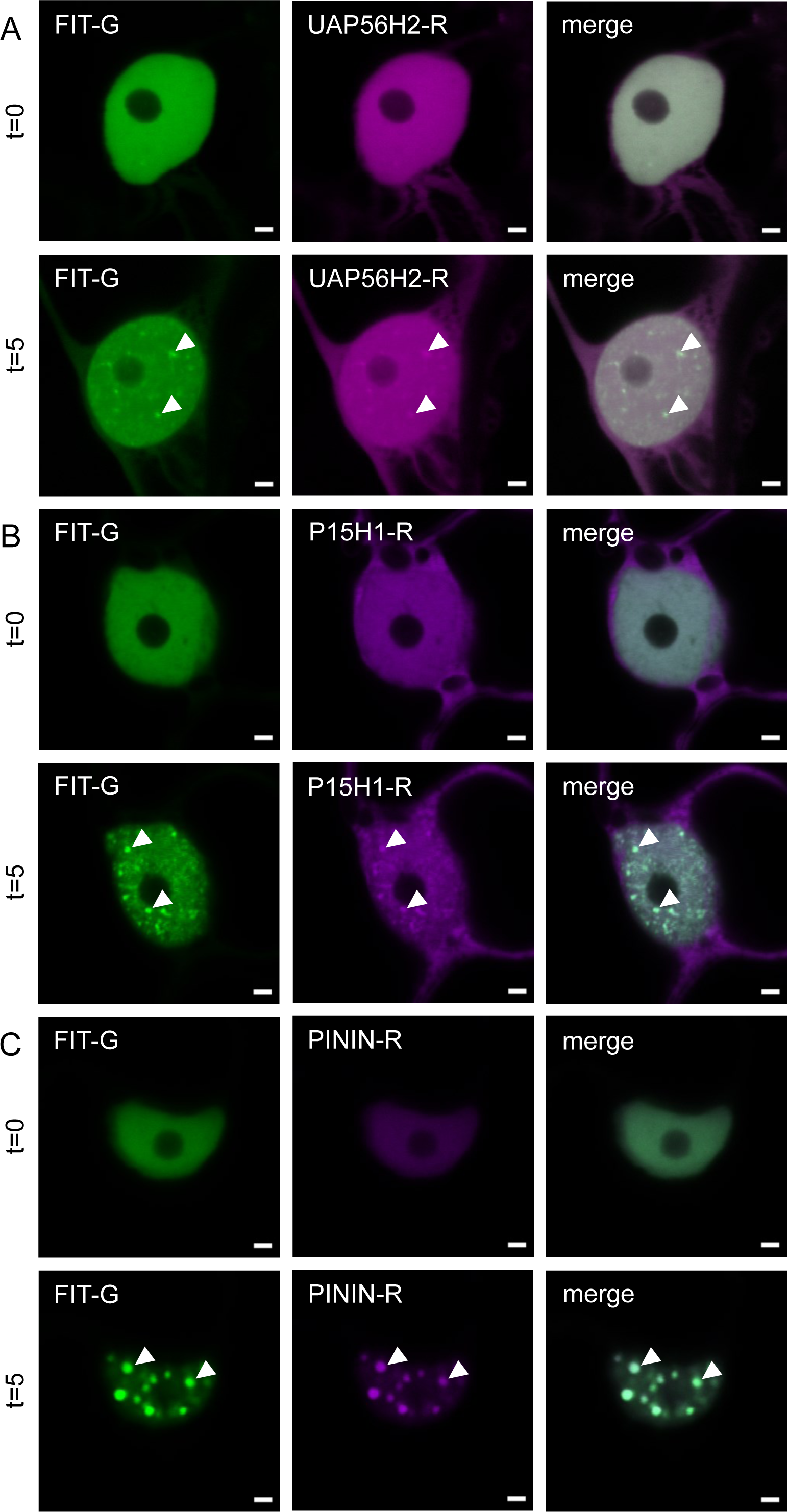
Three NB markers and speckle components became localized in FIT NBs and colocalized fully with FIT (designated type III), suggesting that FIT NBs have speckle function. Confocal images showing localization of FIT-GFP and NB markers (type III) upon co-expression in the nucleus at t=0 and t=5 min. Co-expression of FIT-GFP with A, UAP56H2-mRFP, B, P15H1-mRFP, and C, PININ-mRFP. All three type III NB markers were homogeneously distributed and colocalized with FIT-GFP in the nucleus at t=0, while they colocalized with FIT-GFP in FIT NBs at t=5 min. UAP56H2-mRFP and P15H1-mRFP showed homogeneous localization in the single expression at both t=0 and t=5 min (compare with **Supplemental Figure S4, C and D**), while PININ-mRFP localized mainly in one large and several small NBs upon single expression at t=0 and t=5 min (compare with **Supplemental Figure S4E**). Hence, these three markers adopted the localization of FIT-GFP upon co-expression and suggest that FIT NBs have a speckle function. Scale bar: 2 µm. Arrowheads indicate colocalization within NBs. G = GFP; R = mRFP. Fluorescence protein analysis was conducted in transiently transformed *N. benthamiana* leaf epidermis cells, following the standardized FIT NB analysis procedure. In all examined cells, the proteins showed full colocalization. Representative images from four to seven independent experiments are shown. Contrast of images in A at t=5 was enhanced for better assessment. For data with type I markers (no colocalization) and type II markers (partial colocalization) see **Supplemental Figure S** and **Figure 5**.

Taken together, the colocalization studies underlined the dynamic behavior of inducible FIT NB formation. FIT NBs had a speckle function, in which on the one hand FIT was recruited itself into pre-existing splicing-related NBs (SR45-mRFP and SRm102-mRFP, type II), while on the other hand it also recruited speckle-localized proteins into FIT NBs (UAP56H2-mRFP, P15H1-mRFP, and PININ-mRFP, type III).

### PB components influence FIT NB localization and formation

PBs are plant-specific condensates which harbor various light signaling components (Kircher et al., 2002; Bauer et al., 2004). Among them are the bHLH TFs of the PIF family. As key regulators of photomorphogenesis, they integrate light signals in various developmental and physiological response pathways (Leivar and Monte, 2014; Pham et al., 2018). Indeed, PIF4 may control iron responses in Arabidopsis based on computational analysis of iron deficiency response gene expression networks (Brumbarova and Ivanov, 2019) and both PIF proteins intersect with blue light signaling (Ni et al., 1998; Oh et al., 2004; Pedmale et al., 2016). We tested in the same manner as described above for NB markers, whether FIT NBs coincide with two of the described PB markers, PIF3-mCherry and PIF4-mCherry (Van Buskirk et al., 2014; Qiu et al., 2019, 2021).

We detected distinct localization patterns for PIF3-mCherry and PIF4-mCherry (**Figure 7**). At t=0, PIF3-mCherry was predominantly localized in a single large PB (**Figure 7A**). Localization of single expressed PIF3-mCherry remained unchanged at t=0 and t=15 (**Supplemental Figure S5A**). Upon co-expression, FIT-GFP was initially not present in PIF3-mCherry PB at t=0. After 488 nm excitation, FIT-GFP accumulated and finally colocalized with the large PIF3-mCherry PB at t=15, while the typical FIT NBs did not appear (**Figure 7A**).

**Figure 7:**
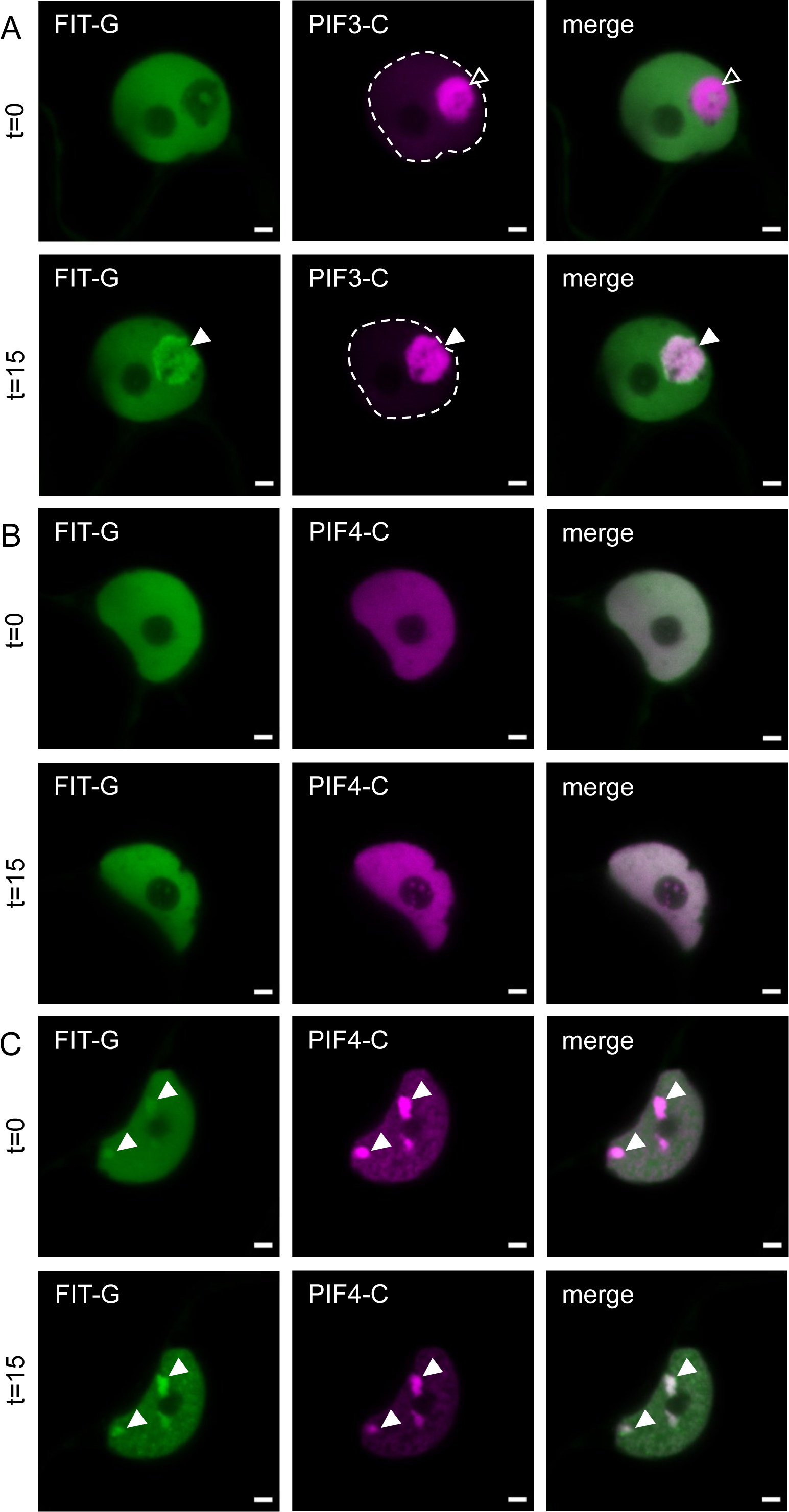
FIT colocalized with photobody (PB) markers in distinct PBs. Confocal images showing localization of FIT-GFP and PB markers upon co-expression in the nucleus at t=0 and t=15 min. Co-expression of FIT-GFP with A, PIF3-mCherry, and B and C, PIF4-mCherry, in B, showing a typical pattern with absence of NBs (ca. 50% of nuclei), in C, showing a typical pattern with presence of NBs (ca. 50% of cells). When FIT-GFP was co-expressed with PB markers, FIT NBs did not appear at t=5 min, but instead, FIT-GFP colocalized with PB markers in PBs at t=15 min. A, PIF3-mCherry localized predominantly to a single large PB at t=0 and t=15 min. FIT-GFP colocalized with PIF3-mCherry in this single large PB at t=15 min. B, PIF4-mCherry and FIT-GFP were both homogeneously distributed in the nucleoplasm at t=0 and t=15 min. In C, FIT-GFP colocalized with PIF4-mCherry in PBs at t=0 and t=15 min. The same localization patterns were found for PIF3-mCherry and PIF4-mCherry upon single expression (compare with **Supplemental Figure S5**). Hence, FIT-GFP was recruited to the two distinct types of PIF3 and PIF4 PBs, whereas PIF3 and PIF4 were not recruited to FIT NBs. This suggests that FIT NBs are affected by the presence of PIF3- and PIF4-containing PBs and a connection to light signaling exists. Scale bar: 2 µm. Arrowheads indicate colocalization in PBs. G = GFP; C = mCherry. Fluorescence protein analysis was conducted in transiently transformed *N. benthamiana* leaf epidermis cells, following the standardized FIT NB analysis procedure. In all examined cells, PIF3 and FIT colocalized fully, while PIF4 and FIT colocalized as indicated in B and C. Representative images from four to six independent experiments are shown.

PIF4-mCherry localized in two different patterns, and both differed substantially from that of PIF3-mCherry. In the one pattern at t=0, PIF4-mCherry was not localized to any PBs, but instead was uniformly distributed in the NP as was the case for FIT-GFP. Such a pattern was also seen at t=15, and then neither PIF4-mCherry nor FIT-GFP were localized in any PBs/NBs (**Figure 7B**). In the other pattern, PIF4-mCherry and FIT-GFP localized in multiple PBs at t=0 and t=15, whereas none of them corresponded morphologically to the typical FIT NBs (**Figure 7C**). The same two localization patterns were also found for PIF4-mCherry in the single expression, whereby 488 nm excitation did not alter PIF4-mCherry localization (**Supplemental Figure S5, B and C**).

Hence, FIT was able to localize to PBs when co-expressed with PIF3 and PIF4, raising the possibility that FIT is a key regulator to cross-connect iron acquisition regulation and light signaling pathways. Again, colocalization demonstrates the dynamic behavior of FIT-GFP.

### Blue light has a promoting effect on iron acquisition responses downstream of FIT

Previous studies have shown that iron uptake by FIT is diurnally and circadian clock-regulated (Vert et al., 2003; Santi and Schmidt, 2009; Salomé et al., 2013). Knowing that FIT NB formation was light-dependent as well as promoting the interaction of the FIT-bHLH039 complex, and that FIT NBs were colocalizing with light signaling and PB components, we reasoned that plants exposed to the FIT NB-forming light cues may show differential FIT-dependent iron uptake responses with respect to control plants exposed to the regular light conditions. To test this, we grew wild-type Arabidopsis plants for 5 d under iron sufficient and iron deficient conditions under white light and exposed the plants additionally for 1 d to blue light. As a control for light quality, we also included exposure to red and far-red light and darkness besides the regular white light. We measured molecular iron uptake responses known to be under the control of FIT-bHLH039 in roots (Gratz et al., 2019).

Iron reductase activity is increased in roots when FIT and bHLH039 are activated in response to iron deficiency in our growth system (Gratz et al., 2019). Iron reductase activity was higher under iron deficient conditions compared to iron sufficient conditions under all white light controls, as expected (**Figure 8A-D**). Interestingly, when seedlings were exposed to blue light for a day, they induced iron reductase activity in the iron deficient versus iron sufficient condition more than in the white light control (**Figure 8A**). On the other hand, exposure to red light did not change the iron reductase activity compared to the white light control (**Figure 8B**), while exposure to far-red light or darkness abolished the induction (**Figure 8, C and D**). Hence, only exposure to blue light had an extra promoting effect on iron uptake in the seedlings compared with the other tested light qualities.

**Figure 8.**
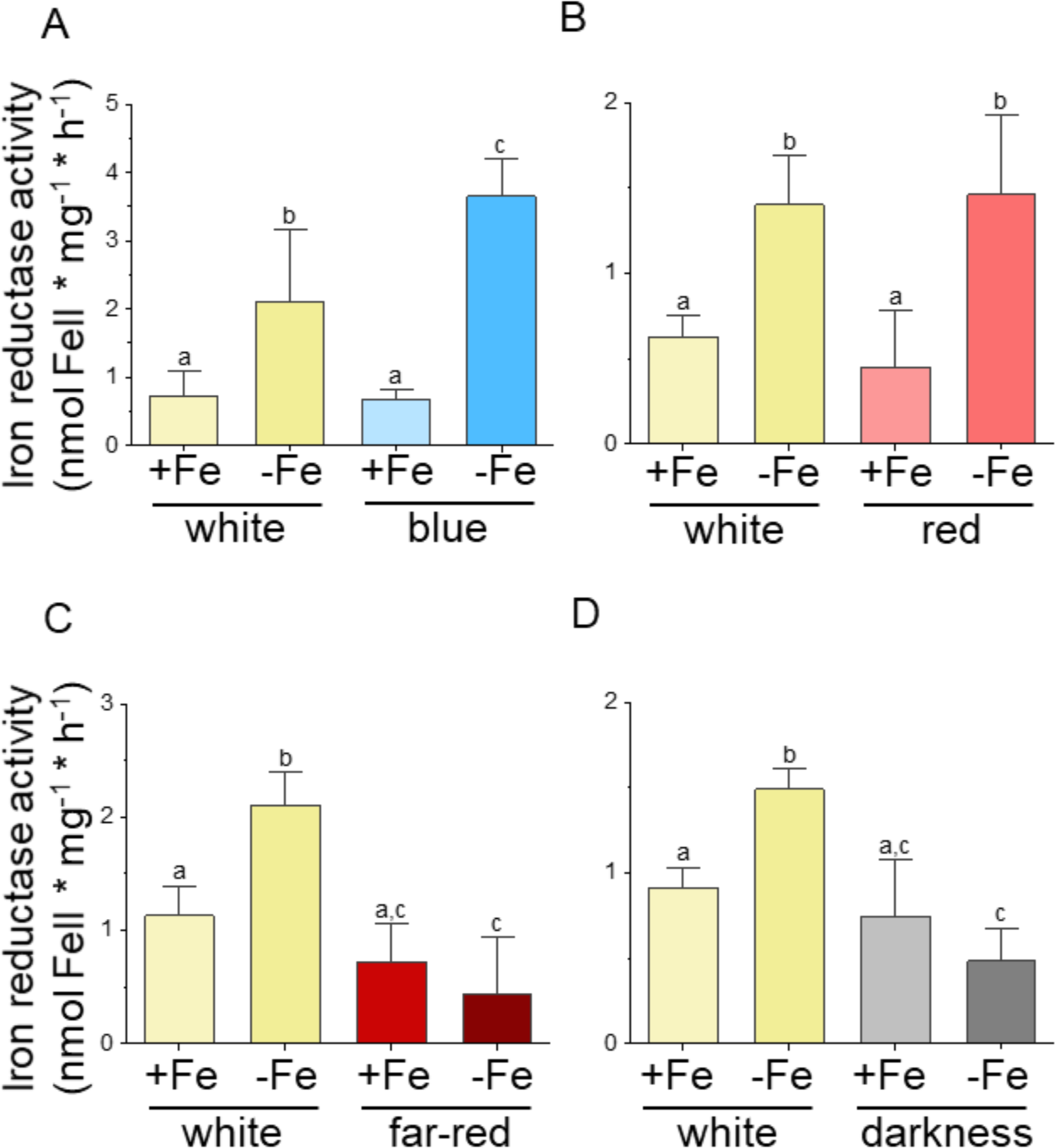
Root iron reductase activity is promoted under blue light. Root iron reductase activity assay on 6-d-old Arabidopsis seedlings grown under white light for 5 d under iron deficient and iron sufficient conditions and then exposed for 1 d to blue, red, far-red light or darkness, or in parallel as control to white light. A, Induction of iron reductase activity upon iron deficiency versus sufficiency was higher under blue light than compared to the white light control. B, Exposure to red light did not change the iron reductase activity compared to the white light control. C-D, Plants exposed to far-red light and darkness did not show induction of iron reductase activity under iron deficient conditions compared to the iron deficient conditions. Two experiments were conducted, one representative result is shown. Bar diagrams represent the mean and standard deviation of four replicates with four seedlings each (n=4). Statistical analysis was performed with one-way ANOVA and Tukey post-hoc test. Different letters indicate statistically significant differences (P < 0.05).

Increased iron acquisition under iron deficiency and enhanced iron reductase activity require FERRIC REDUCTASE-OXIDASE2 (FRO2) protein (Yi and Guerinot, 1996; Robinson et al., 1999), and *FRO2* to be induced, along with *IRON REGULATED TRANSPORTER1* (*IRT1*; Eide et al., 1996; Korshunova et al., 1999). *FRO2* and *IRT1* are target genes of FIT-bHLH039 in our growth system (Gratz et al., 2019, 2020). As a readout of the activity of the FIT-bHLH039 complex, we examined gene regulation of *FRO2* and *IRT1* as well as *FIT* and *BHLH039* under the same light conditions as described above. Previously, it was shown that the expression of *IRT1* and *FRO2* was downregulated under iron deficient conditions in darkness compared to iron deficient conditions in white light, while the opposite was true for *FIT* (Santi and Schmidt, 2009).

Under white light conditions, gene expression for all four genes was tested in four independent samples. In 15 out of the 16 cases, gene expression of *FIT*, *BHLH039*, *FRO2* and *IRT1* was significantly higher under iron deficient than sufficient conditions, as expected, and in only one case for *FIT*, it was similar and not significantly enhanced (**Figure 9A-P, with exception of E**, *FIT*). Under blue light, *FIT* and *BHLH039* expression was induced upon iron deficiency in the same manner as in white light (**Figure 9, A and B**), whereas *FRO2* and *IRT1* were induced to a higher level in response to iron deficiency in blue light versus white light (**Figure 9A-D**). Again, plant responses were different upon exposure to other light conditions. *FIT* and *IRT1* were similarly induced, while *BHLH039* and *FRO2* were even less induced by iron deficiency under red light than white light (**Figure 9E-H**). In both, the far-red light and darkness situations, *FIT* was induced under iron deficiency versus sufficiency, while on the other side, *BHLH039*, *FRO2* and *IRT1* were not induced at all in these light conditions (**Figure 9I-P**). *FIT* was similarly induced under far-red light and white light (**Figure 9I**), whereas in darkness it was even induced to higher level (**Figure 9M**), as previously reported (Santi and Schmidt, 2009).

**Figure 9.**
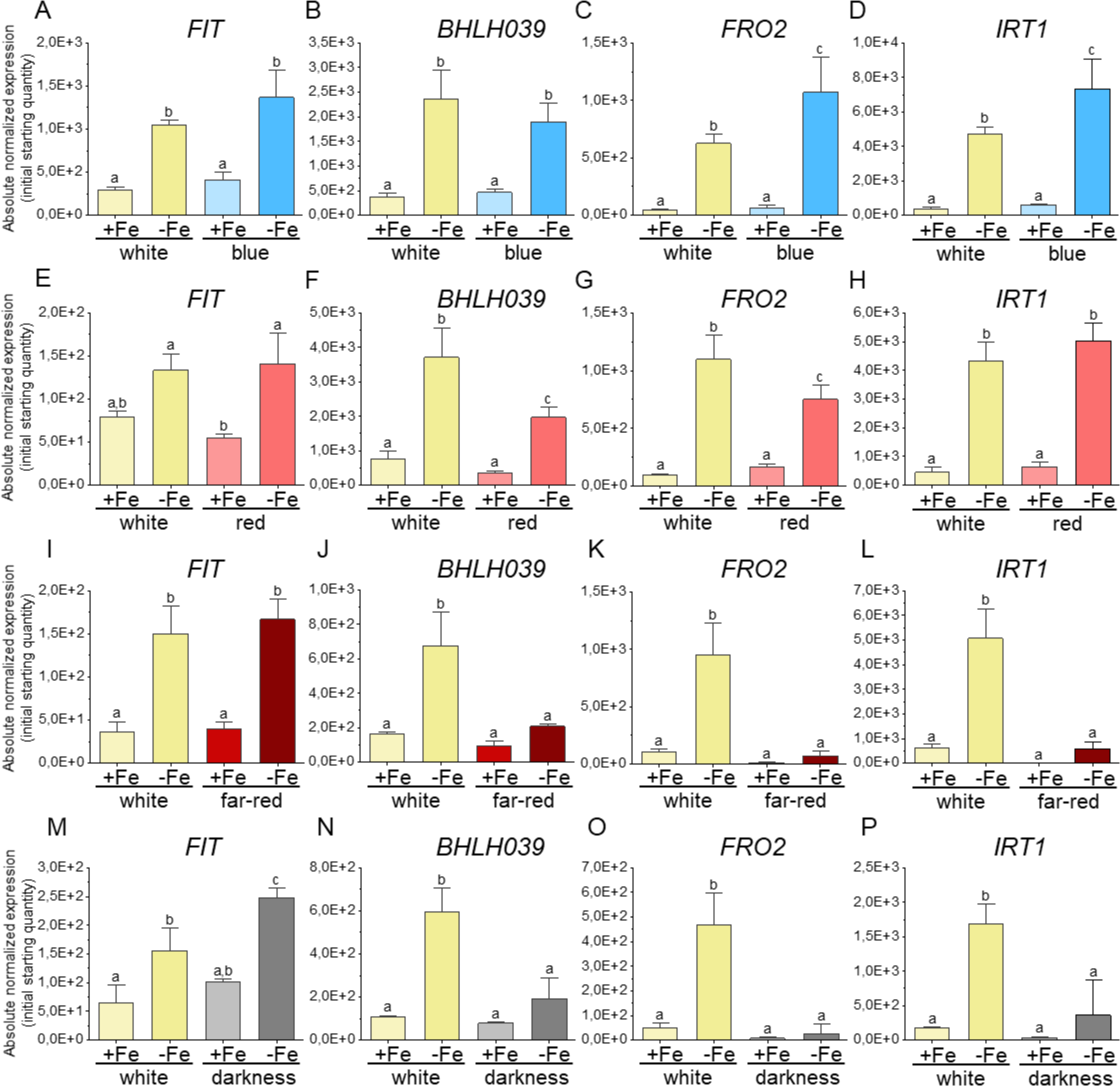
Induction of FIT target gene expression is enhanced under blue light. Gene expression analysis of iron deficiency genes *FIT*, *BHLH039*, *FRO2*, and *IRT1* on 6-d-old Arabidopsis seedlings grown under white light for 5 d under iron deficient and iron sufficient conditions and then exposed for 1 d to blue, red, far-red light or darkness, or in parallel as control to white light. A-B, *FIT* and *bHLH039* gene induction in response to low iron supply did not change after blue light exposure compared to the white light control. C-D, *FRO2* and *IRT1* gene expression increased after blue light exposure in iron deficient conditions compared to respective iron sufficient condition and white light control. E, *FIT* gene expression did not change after red light exposure compared to the white light control. F-G, *BHLH039* and *FRO2* gene expression decreased after red light exposure in iron deficient conditions compared to respective iron sufficient condition and white light control. H, *IRT1* gene expression did not change after red light exposure compared to the white light control. I, *FIT* gene expression did not change after far-red light exposure compared to the white light control. J-L, *BHLH039*, *FRO2*, and *IRT1* gene expression decreased after far-red light exposure in iron deficient conditions compared to white light deficient condition, comparable to the respective iron sufficient condition and iron sufficient white light control. M, *FIT* gene expression was higher in darkness under iron deficient conditions compared to respective iron sufficient condition and white light control. N-P, *BHLH039*, *FRO2*, and *IRT1* gene expression decreased after being exposed to darkness in iron deficient conditions compared to white light deficient condition, comparable to the respective iron sufficient condition and iron sufficient white light control. Two experiments were conducted, one representative result is shown. Bar diagrams represent the mean and standard deviation of three replicates with twelve seedlings and two technical replicates each (n=3). Statistical analysis was performed with one-way ANOVA and Tukey post-hoc test. Different letters indicate statistically significant differences (P < 0.05).

Since colocalization with NB markers showed partial or full colocalization with FIT within NBs (**Figure 5 and 6**), we included gene expression analysis on some of the previously described intron retention (IR) splicing variants of *FIT*, *BHLH039*, and FIT target genes *IRT1* and *FRO2* (Li et al., 2013, 2016a) to see whether differential alternative splicing would be detectable under the same experimental conditions that led to FIT NB formation in Arabidopsis (**Figure 1A**). For this, we grew wild-type Arabidopsis plants under iron sufficient and iron deficient conditions for 5 d under white light and exposed the plants for 1.5 or 2 h to blue light.

IR splicing variants occurred to higher extent under iron deficient conditions compared to iron sufficient conditions regardless of the light condition (**Supplemental Figure S6**). After normalization of gene expression of plants grown under iron deficient conditions with plants grown under iron sufficient conditions, no difference between the respective gene and its splicing variants could be shown (**Supplemental Figure S7**).

To sum up, the FIT-bHLH039-dependent target gene expression is similarly light quality-dependent as the FIT-bHLH039-dependent iron reductase activity but not the splicing variant abundance of the tested genes. Hence, very interestingly, blue light cues which lead to FIT NB formation, promoted FIT-bHLH039-dependent iron uptake responses, fitting with the above finding that FIT NBs are linked with active TF complexes in condensates. However, a differential effect of NB-forming conditions and alternative splicing was not found suggesting that FIT NBs do not primarily affect alternative splicing processes of the tested iron deficiency marker genes.

## Discussion

In this study, we uncovered a previously unknown phenomenon, the blue light-induced and reversible accumulation of FIT condensates in FIT NBs. LLPS was most likely the underlying mechanism for this highly dynamic process. FIT NBs were enriched in active FIT TF complexes that are required for iron deficiency gene regulation. FIT associated with speckles and PBs in a highly dynamic fashion, indicating a function in transcriptional and posttranscriptional control. Blue light exposure resulted in enhanced molecular FIT-bHLH039-dependent iron acquisition responses but not a different effect on alternative splicing pathways of *FIT*, *BHLH039*, *IRT1* and *FRO2*. Based on these data, we propose that FIT NBs are dynamic microenvironments with active FIT TF complexes that possibly are hubs to enhance and cross-connect transcriptional iron deficiency gene expression with posttranscriptional regulation and environmental signaling.

### A standardized procedure for FIT NB induction was crucial to delineate the characteristics of FIT NBs in reliable manner

We were able to characterize the nature and potential function of light-induced FIT NBs using quantitative microscopy-based techniques and a standardized NB analysis procedure. Roots also formed NBs, however their direct inspection was not possible using the same microscopy techniques. Since condensation depleted FIT protein in the nucleoplasm, the remaining low FIT protein concentration can be the reason why FIT NBs remained few in number in the Arabidopsis root cells. The *N. benthamiana* protein expression system did not present these limitations and high-quality measurement data were obtained for all experimental series. Furthermore, this expression system is a well-established and widely utilized system in plant biology (Martin et al., 2009; Bleckmann et al., 2010; Leonelli et al., 2016; Burkart et al., 2022). The developed standardized assay generated reliable and accurate data for statistical analysis and quantification to conclude about FIT NB characteristics.

Condensation likely explains the reduced mobility of FIT-GFP versus FITmSS271AA-GFP seen in a previous study (Gratz et al., 2019). In liquid state, condensates are more mobile than in the solid one. The condensate formation was reversible, speaking in favor of a liquid state of the condensates. According to FRAP data, FIT NBs maintained a dynamic exchange of FIT protein with the surrounding NP. FIT NBs were also mostly of circular shape. Circular condensates appear as droplets, in contrast to solid-like condensates that are irregularly shaped (Shin et al., 2017). These characteristics speak in favor of liquid-like features, suggesting that LLPS underlies FIT NB formation. A similar situation was described for CRY2 PBs, which were also reversible and of circular shape with mobile protein inside PBs (Wang et al., 2021). bHLH039 was found accumulated in cytoplasmic foci at the cell periphery (Trofimov et al., 2019). In such foci, bHLH039 was immobile, and we suspect it was in a non-functional state in the absence of FIT. This underlines the understanding that liquid condensates such as FIT NBs are dynamic microenvironments, whereas immobile condensates point rather towards a solid and pathological state (Shin et al., 2017).

Additionally, the partial and full colocalization of FIT NBs with various previously reported NB markers confirm that FIT indeed accumulates in and forms NBs. Since several of NB body markers are also behaving in a dynamic manner, this corroborates the formation of dynamic FIT NBs affected by environmental signals.

In conclusion, the properties of liquid condensation and colocalization with NB markers, along with the findings that it occurred irrespective of the fluorescence protein tag preferentially with wild-type FIT, allowed us to coin the term of ‘FIT NBs’.

### IDR^Ser271/272^ was crucial for interaction and NB formation of FIT

FIT NBs were hotspots for homodimeric and heterodimeric FIT interaction, allowing to assume that they are integrated in the iron deficiency response as interaction hubs. These abilities distinguished wild-type FIT and mutant FITmSS271AA, indicating that wild-type FIT is a multivalent protein and that IDR^Ser271/272^ is important for that. FIT protein may interact via the helix-loop-helix interface and via its C-terminus containing the IDR (Lingam et al., 2011; Le et al., 2016; Gratz et al., 2019) to be multivalent. The flexible IDRs adapt to multiple protein partners and are therefore crucial for multivalency (Tarczewska and Greb-Markiewicz, 2019; Emenecker et al., 2020; Salladini et al., 2020), and the amino acid composition of IDRs is important for condensation (Powers et al., 2019; Emenecker et al., 2021; Huang et al., 2022). Very interestingly, posttranslational modification in form of phosphorylation within IDRs may regulate condensate formation (Owen and Shewmaker, 2019). Ser271/272 is targeted by a FIT-interacting protein kinase that can affect FIT activity and FIT phosphorylation (Gratz et al., 2019). Hence, phosphorylation of Ser271/272 might perhaps be a trigger for NB formation *in vivo*. It could not be distinguished in this study whether FIT homodimers were a prerequisite for the localization of bHLH039 in NBs or whether FIT-bHLH039 complexes also initiated NBs on their own.

### Formation of FIT NB is highly dynamic

FIT may have formed FIT NBs as entirely newly formed structures upon the light trigger. But it is also possible that FIT joined pre-existing NBs, which then became the structures we termed FIT NBs. Partial or full colocalization of FIT-GFP with NB and PB markers revealed the remarkably high and intriguing dynamic nature of FIT NBs and suggests that both possibilities are plausible. FIT NBs are light-triggered, and this speaks in favor of pre-existing NBs. Since FIT does not possess light-responsive domains, it is most likely that a light-responsive protein must be inducing FIT NB formation. The basic leucine zipper TF ELONGATED HYPOCOTYL5 (HY5) could be a good candidate, since HY5 is a mobile protein involved in iron acquisition in tomato (Gao et al., 2021; Guo et al., 2021) and acts positively on iron uptake and gene expression of *FIT*, *FRO2*, and *IRT1* in Arabidopsis (Mankotia et al., 2023). Possibly activation and condensation involve not only the studied NB and PB markers but also potentially signaling proteins or further scaffold proteins that are part of the multivalent protein complexes in FIT NBs. On the other hand, FIT-GFP accumulated not only in FIT NBs but also in the pre-existing NBs with type II NB markers (SR45 and SRm102) after the FIT NB induction. In this respect, type II markers were similar to PIF3 and PIF4. FIT-GFP was recruited to pre-existing PBs and again only after the light trigger. Interestingly, typical FIT NB formation did not occur in the presence of PB markers, indicating that they must have had a strong effect on recruiting FIT. This is interesting because the partially colocalizing SR45, PIF3 and PIF4 are also dynamic NB components. Active transcription processes and environmental stimuli affect the sizes and numbers of SR45 speckles and PB (Ali et al., 2003; Legris et al., 2016; Meyer, 2020). This may indicate that, similarly, environmental signals might have affected the colocalization with FIT and resulting NB structures in our experiments. Another factor of interference might also be the level of expression. Overall, the dynamics of FIT colocalization with type II NB and PB markers suggest that these condensates dictated FIT condensation in their own pre-existing NBs/PBs. This recruiting process could be navigated via protein-protein interaction since this is the driving force of condensation (Kaiserli et al., 2015; Emenecker et al., 2020).

On the other side, the full colocalization of FIT with type III NB markers speaks in favor of a *de novo* FIT NB formation. The three fully colocalizing type III NB markers (UAP56H2, P15H1 and PININ) accumulated only in FIT NBs upon co-expression with FIT and mostly not on their own. The same was true for bHLH039, that joined FIT in FIT NBs, showing that FIT not only facilitated bHLH039 nuclear localization (Trofimov et al., 2019) but also condensation. Interestingly, FIT was able to change PININ nuclear localization. In single expression, PININ was localized to a major large NB, but in colocalization with FIT it joined the typical FIT NBs. This suggests that FIT dictates bHLH039 and type III NB markers and highlights that FIT is also able to set the tone for NB formation. Hence, FIT can recruit other proteins into NBs, and it is possible that FIT forms its own NBs. Protein-protein interaction could underly this recruitment, as evident for bHLH039 (Kaiserli et al., 2015; Emenecker et al., 2020).

### FIT NBs have speckle characteristics

Since the type II and III markers are splicing components, the colocalization studies suggest that FIT NBs are speckles, in agreement with the dynamic nature of FIT NBs. Speckles are highly dynamic, forming around transcriptionally active sites in the interchromatin regions recruiting several protein functions like mRNA synthesis, maturation, splicing and export (Reddy et al., 2012; Galganski et al., 2017). Co-transcriptional splicing in plants is also recently rising (Nojima et al., 2015; Zhu et al., 2018; Chaudhary et al., 2019). Mediator complex condensation was shown to drive transcriptional control (Boija et al., 2018) and interestingly, FIT was also shown to interact with Mediator complexes, directly and indirectly (Yang et al., 2014; Zhang et al., 2014). Besides, other studies suggest TF condensation to be involved in transcriptional regulation (Kaiserli et al., 2015; Boija et al., 2018; Brodsky et al., 2020; Huang et al., 2022). The type II speckle component SR45, is a highly mobile protein in speckles and required phosphorylation for proper speckle localization (Ali et al., 2003; Reddy et al., 2012). These processes fit well to the described FIT NB attributes. The characterization of FIT NBs as speckles is interesting because regulation of splicing and epigenetic regulation is associated with both, iron deficiency gene expression and SR45 (Fanara et al., 2022; Mikulski et al., 2022). UAP56H2, P15H1, and PININ (type III) are connected to SR45 and SRm102 (type II) in mammalian cells as all being part of the exon junction complex and interacting with each other (Lin et al., 2004; Pendle et al., 2005). This is an interesting parallel, as it suggests that type II and type III marker localization is conserved across kingdoms, underlying the ancient nature of condensates. Indeed, SR45 and PININ located both to a very large NB in non-induced cells. This opens the possibility that the two proteins might localize to the same type of speckle, as also might FIT. Taken together, the observations confirm the high diversification and complexity of FIT NBs and speckles (Lorković et al., 2008) and it is tempting to speculate that FIT might regulate splicing and alternative splicing of its target genes by recruiting speckle components.

However, further studies are needed to investigate whether the blue light treatment resulting in FIT NBs causes alternative splicing compared with white light. The previously reported intron retention events (Li et al., 2013) were confirmed in our study but a differential effect of light was not apparent, although it may be rare and might have been masked by non-reactive nuclear events.

### Model and physiological integration of FIT NBs

FIT NBs may serve the rapid rearrangement of TFs to enhance target gene expression and subsequent physiological responses during a high photosynthetic period. The formation and dissociation of NBs could be a fast way to adjust iron uptake and specifically uncouple *IRT1* and *FRO2* from FIT regulation and therefore adapt even better to environmental changes, possibly triggered by phosphorylation (Gratz et al., 2020). The rapid speed by which FIT NB appeared within 5 min in *N. benthamiana* leaf cells speaks in favor of protein rearrangement rather than protein synthesis. The long duration of FIT NB formation after blue light induction in Arabidopsis roots suggests that signal transduction was more complex and possibly involved intracellular or even cell-to-cell and long-distance leaf-to-root signaling. A long-distance signal or a signaling cascade triggered by light might be involved in FIT NB formation. CRY1/CRY2 and HY5 are promising candidates for further studies (Gao et al., 2021; Guo et al., 2021; Mankotia et al., 2023). In order to undergo phase separation, a certain protein concentration must be reached (Bracha et al., 2018). Since FIT protein is subject of proteasomal turnover in roots, FIT NB formation may depend on FIT protein interaction partners in roots that need to be activated (Lingam et al., 2011; Meiser et al., 2011).

In summary, FIT localizes to dynamic and reversible NBs. FIT NBs contain active TF complexes for iron acquisition gene expression and are speckles that link transcriptional with posttranscriptional regulation. The appearance of FIT NBs is inducible by blue light, a condition that promotes iron mobilization responses, and light regulated PB components are connected with FIT NBs and vice versa (**Figure 10**). It will be interesting in the future to test hormonal and environmental triggers that may stabilize FIT protein prior to examining the initiation of FIT NBs in root physiological situations. Further studies are needed to determine whether FIT NBs might have a transcriptional and posttranscriptional function to regulate unknown FIT target sites.

**Figure 10:**
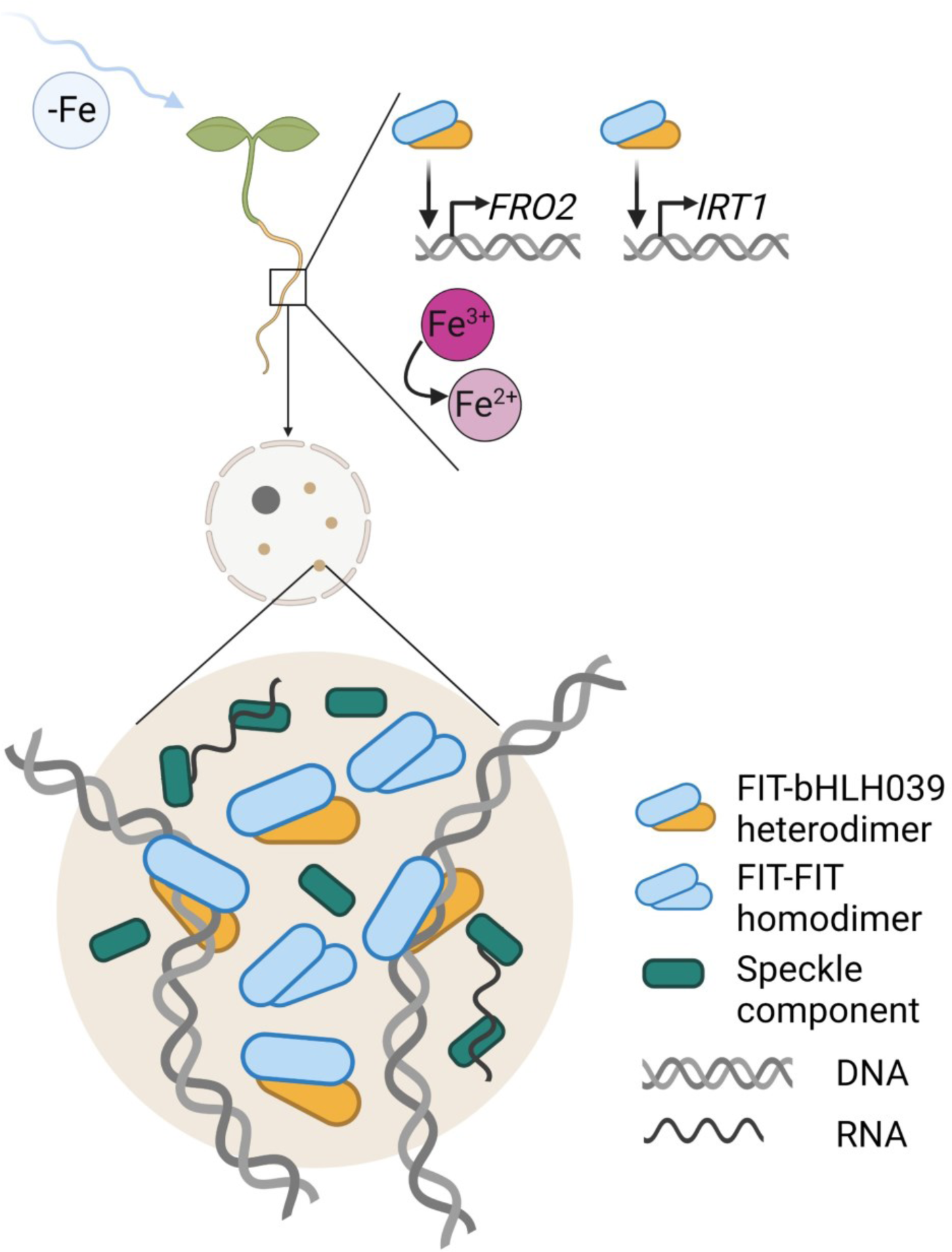
Summary model illustrating the effect of blue light on FIT NB formation and iron uptake, suggesting that FIT NBs are related to transcriptional and posttranscriptional regulation in speckles and that blue light has a promoting effect on iron uptake. FIT accumulates in FIT NBs upon induction with blue light. FIT NBs are reversible, dynamic and of circular shape and may undergo LLPS. FIT homodimers and FIT-bHLH039 heterodimers are present in FIT NBs. FIT-bHLH039 is an active TF complex for upregulating the expression of iron acquisition genes in roots (Gratz et al., 2019). Hence, FIT NBs are subnuclear sites related to transcriptional regulation and because of their colocalization with speckle components, also to speckles. Blue light enhances root iron reductase activity and gene expression of *FRO2* and *IRT1*, all of which are downstream of FIT-bHLH039. In summary, FIT NBs are blue light-inducible subnuclear sites where active TF complexes of FIT and bHLH039 accumulate, linking transcriptional and posttranscriptional regulation in speckles with a promoting blue light effect on iron uptake.

## Materials and methods

### Plant material and growth conditions

*Arabidopsis thaliana* experiments were conducted with the Col-0 accession. Arabidopsis line FIT_pro_:FIT-GFP used for imaging was obtained as follows: First, the FIT_pro_:FIT-GFP construct was obtained in a plant gene expression binary vector via GreenGate cloning (Lampropoulos et al., 2013). *FIT* promoter sequence of 2012 bp upstream of the CDS was cloned into pGGA000 entry vector using pFIT GG fw and pFIT GG rv primers with respective overhang sequences (**Supplemental Table S1**). FIT CDS (954 bp, without stop codon) was cloned into pGGC000 entry vector using cFIT GG fw and cFIT GG rv primers with respective overhang sequences (**Supplemental Table S1**). The final construct was created via the GreenGate reaction with the pGGZ001 destination vector, containing the entry vectors pGGA000 FIT promoter, pGGB003 *B-dummy* for N-tag, pGGA000 FIT CDS, pGGD001 *linker*-GFP, pGGE009 *UBQ10* terminator, and pGGF001 *p^d^MAS^e^:BastaR^f^:t^g^MAS*, and verified by sequencing. The plasmid was transferred to *Rhizobium radiobacter* pSOUP strain (GV3103). Transgenic plants (in Col-0 wild-type background) were obtained by floral dip transformation (Clough and Bent, 1998) and subsequent Basta selection. *Arabidopsis thaliana* 2x35S_pro_:FIT-GFP/*fit-3* line used for imaging was previously described in Gratz et al. (2019).

*Arabidopsis thaliana* FIT_pro_:FIT-GFP and 2x35S_pro_:FIT-GFP/*fit-3* seedlings were used for localization studies. Seeds were sterilized and grown upright on Hoagland medium plates (macronutrients: 1.5 mM Ca(NO_3_)_2_ · 4H_2_O, 0.5 mM KH_2_PO_4_, 1.25 mM KNO_3_, 0.75 mM MgSO_4_ · 7H_2_O; micronutrients: 0.075 µM (NH_4_)6Mo_7_O_24_ · 4H_2_O, 1.5 µM CuSO_4_ · 5H_2_O, 50 µM H_3_BO_3_, 50 µM KCl, 10 µM MnSO_4_ · H_2_O, 2 µM ZnSO_4_ · 7H_2_O; 1.4 % (w/v) plant agar, 1 % (w/v) sucrose, pH 5.8) with no iron supply for 5 d under long day conditions (16 h light/8 h dark) at 21°C in a plant chamber (CLF Plant Climatics) under white light (120 µmol m^-2^ s^-1^).

For iron reductase activity assays and gene expression analysis, *Arabidopsis thaliana* seedlings were sterilized and grown upright on Hoagland medium plates with either no iron supply or with 50 µM FeNaEDTA supply for 5 d under long day conditions (16 h light/8 h dark) at 21°C in a plant chamber (CLF Plant Climatics) under white light (120 µmol m^-2^ s^-1^). For different light treatments plates were transferred to LED light chambers (CLF Plant Climatics) for 1.5 h-2 h (splicing variant gene expression analysis) or 1 d under continuous blue light (70 µmol m^-2^ s^-1^), continuous red light (55 µmol m^-2^ s^-1^), continuous far-red light (65 µmol m^-2^ s^-1^) or darkness. White light was used as control (125 µmol m^-2^ s^-1^).

*Nicotiana benthamiana* plants were grown in the greenhouse facility for approx. 4 weeks under long day conditions (16 h light/8 h dark) for use in transient protein expression experiments.

### Root iron reductase activity assay

6-d-old seedlings were grown as described in the text. Root iron reductase activity was determined as described in Le *et al*. (2016). Plants were washed with 0.1 M Ca(NO_3_)_2_ · 4H_2_O and subsequently incubated for 1 h in darkness at room temperature in the iron reductase solution (300 µM Ferrozine, 100 µM FeNaEDTA). The absorbance of the ferrozine-Fe^2+^ complex was measured at 562 nm and used to calculate the root iron reductase activity normalized to root fresh weight.

### Gene expression analysis by RT-qPCR

5-d or 6-d-old seedlings were grown as described in the text. The reverse transcription-real time-quantitative polymerase chain reaction (RT-qPCR) procedure and analysis is described in Abdallah and Bauer (2016). Briefly, plant material was harvested, using three biological replicates, and shock frozen. RNA was extracted (peqGOLD Plant RNA Kit, PEQLAB). Subsequently, an amount of 0.5 µg of total RNA was used for cDNA synthesis (Thermo Scientific). Diluted cDNA, SYBR green master mix (Thermo Scientific) and respective primers (**Supplemental Table S1**) were pipetted in a 96 well plate and qPCR was performed with two technical duplicates with the CFX96 Touch^TM^ Real-Time PCR Detection System (Bio-Rad). qPCR data were analyzed, and quantification was obtained according to the mass standard curve analysis procedure. Normalization was performed using the reference gene *ELONGATION FACTOR1Α* (*EFc*). Statistical analysis was performed with absolute normalized gene expression levels.

### Microscopy of *Arabidopsis thaliana* seedlings

Protein localization studies in roots of 5-d-old seedlings of the *Arabidopsis thaliana* line FIT_pro_:FIT-GFP and 2x35S_pro_:FIT-GFP/*fit-3* were performed with the widefield microscope ELYRA PS (Zeiss) equipped with a EMCCD camera. To induce FIT NB formation, whole seedlings were exposed to 488 nm laser light for several minutes. GFP was excited with a 488 nm laser and detected with a BP 495-575 + LP 750 beam splitter. Images were acquired with the C-Apochromat 63x/1.2 W Korr M27 (Zeiss) objective, pixel dwell time of 1.6 µs and frame size of 512x512. Pictures were processed with the manufacturer’s software ZEN lite (Zeiss).

### Generation of fluorescent constructs

All constructs used in this study are listed in **Supplemental Table S2**. Generation of fluorescent translational C-terminal fusion of PIF3 and PIF4 with mCherry was performed with Gateway Cloning. CDS of *PIF3* was amplified with the PIF3 GW fw and PIF3 GW rv primers (**Supplemental Table S1**), and CDS of *PIF4* was amplified with the PIF4 GW fw and PIF4 GW rv primers (**Supplemental Table S1**), and introduced into the entry vector pDONR207 via the BP reaction (Life Technologies) and subsequently into the inducible pABind 35S_pro_:mCherry destination vector (Bleckmann et al., 2010) via the LR reaction (Life Technologies). Finally, *Rhizobium radiobacter* was transformed with the constructs for transient transformation of *Nicotiana benthamiana* leaf epidermal cells.

### Transient transformation of *Nicotiana benthamiana* leaf epidermal cells

Transient protein expression was performed in *Nicotiana benthamiana* leaf epidermal cells according to Bleckmann et al. (2010). This was performed for localization studies, FRAP measurements, anisotropy (homo-FRET) measurements, FRET-FLIM measurements, and NB quantification. Cultures of *Rhizobium radiobacter* containing the construct of interest (**Supplemental Table S2**) were incubated overnight and cell were pelleted and dissolved in AS medium (250 µM acetosyringone (in DMSO), 5 % (w/v) sucrose, 0.01 % (v/v) silwet, 0.01 % (w/v) glucose). An OD_600nm_ of 0.4 was set for all constructs. A *Rhizobium radiobacter* strain containing the silencing repressor p19 vector (Shamloul et al., 2014) was used additionally for bHLH039-mCherry to enhance expression. After 1 h incubation on ice the suspension was infiltrated with a syringe into the abaxial side of the leaf. *Nicotiana benthamiana* plants were kept under long day conditions (16 h light/8 h dark) in the laboratory after infiltration. Imaging was performed 2-3 d after infiltration. Expression of constructs with an inducible 35S promoter was induced 16 h prior to imaging with β-estradiol (20 µM β-estradiol (in DMSO), 0.1 % (v/v) Tween 20). In total, 3-4 differently aged leaves of 2 plants were infiltrated and used for imaging. One infiltrated leaf with homogenous presence of one or two fluorescence proteins was selected, depending on the aim of the experiment, and ca. 30 cells were observed. Images are taken from 3-4 cells, one representative image is shown.

### Confocal microscopy

For localization studies a confocal laser scanning microscope LSM780 (Zeiss) was used. Imaging was controlled by the ZEN 2.3 SP1 FP3 (Black) (Zeiss) software. GFP was excited with a 488 nm laser and detected in the range of 491-553 nm. mCherry and mRFP were excited with a 561 nm laser and detected in the range of 562-626 nm. Fluorophore crosstalk was minimized by splitting of the excitation tracks and reduction of emission spectrum overlap. Images were acquired with the C-Apochromat 40x/1.20 W Korr M27 (Zeiss) objective, zoom factor of 8, pinhole set to 1,00 AU, pixel dwell time of 1.27 µs and frame size of 1.024x1.024. Z-stacks for quantification were taken with the same settings, except with pixel dwell time of 0.79 µs and frame size of 512x512. Pictures were processed with the manufacturer’s software ZEN lite (Zeiss).

### Standardized FIT NB analysis procedure

Following *Nicotiana benthamiana* leaf infiltration with *Rhizobium radiobacter*, FIT-GFP protein expression was induced after 2-3 d by β-estradiol, as described above. 16 h later, a leaf disc was excised and FIT-GFP fluorescence signals were recorded (t=0). The leaf disc was excited with 488 nm laser light for 1 min. 5 min later, FIT-GFP accumulation in FIT NBs was observed (t= 5 min). See **Supplemental Figure S1C**. This procedure was modified by using different time points for NB analysis and different constructs (**Supplemental Table S2**) and co-expression as indicated in the text. Imaging was performed at the respective wavelengths for detection of GFP and mRFP/mCherry.

### FRAP measurements

FRAP measurements (Bancaud et al., 2010; Trofimov et al., 2019) were performed at the confocal laser scanning microscope LSM780 (Zeiss). Imaging was controlled by the ZEN 2.3 SP1 FP3 (Black) (Zeiss) software. GFP was excited with a 488 nm laser and detected in the range of 491-553 nm. Images were acquired with the C-Apochromat 40x/1.20 W Korr M27 (Zeiss) objective, zoom factor of 8, pinhole set to 2,43 AU, pixel dwell time of 1.0 µs, frame size of 256x256, and 300 frames. After 20 frames, a NB was bleached with 50 iterations and 100% 488 nm laser power. Fluorescence intensity was recorded for the bleached NB (ROI), a non-bleached region equal in size to the NB (BG) as well as for the total image (Tot). Values were calculated and processed in Excel (Microsoft Corporation). Background subtraction and normalization to calculate the relative fluorescence intensity was performed as follows: [(ROI(t)-BG(t)/Tot(T)-BG(t))*(Tot(t0)-BG(t0)/ROI(t0)-BG(t0))]. The mobile fraction was calculated as follows: [(F_end_-F_post_)/(F_pre_-F_post_)*100]. F_pre_ marks the average of the 20 values before bleaching, F_post_ marks the value right after the bleaching, and F_end_ marks the average of the 280 values after the bleaching. Pictures were processed with the manufacturer’s software ZEN lite (Zeiss).

### Anisotropy (homo-FRET) measurements

Anisotropy measurements (Stahl et al., 2013; Weidtkamp-Peters et al., 2022) were performed at the confocal laser scanning microscope LSM780 (Zeiss) equipped with a polarization beam splitter, bandpass filter (520/35), and a single-photon counting device HydraHarp (PicoQuant) with avalanche photo diodes (τ-SPADs). Emission was detected in parallel and perpendicular orientation. Rhodamine 110 was used to determine the G factor to correct for the differential parallel and perpendicular detector sensitivity. Calibration of the system was performed for every experiment and measurements were conducted in darkness. Free GFP and GFP-GFP were used as references for mono- and dimerization, respectively. GFP was excited with a linearly polarized pulsed (32 MHz) 485 nm laser and 0.05-1 µW output power. Measurements were recorded with a C-Apochromat 40x/1.20 W Korr M27 (Zeiss) objective, zoom factor of 8, pixel dwell time of 12.5 µs, objective frame size of 256x256, and 40 frames. Measurements were controlled with the manufacturer’s ZEN 2.3 SP1 FP3 (Black) (Zeiss) software and SymPhoTime 64 (PicoQuant) software. SymPhoTime 64 (PicoQuant) software was used for analysis in the respective regions of interest (whole nucleus, NB, NP) and to generate color-coded FA value images. Minimal photon count was set to 200.

### FRET-FLIM measurements

FRET-FLIM measurements (Borst and Visser, 2010; Weidtkamp-Peters and Stahl, 2017) were taken at the confocal laser scanning microscope FV3000 (Olympus) equipped with a multi-photon counting device MultiHarp 150 (PicoQuant) with avalanche photo diodes (τ-SPADs) and bandpass filter (520/35). Erythrosine B (quenched in saturated potassium iodide) was used to record the Instrument Response Function to correct for the time between laser pulse and detection. Calibration of the system was performed for every experiment and measurements were conducted in darkness. FIT-GFP and FITmSS271AA-GFP were used as negative controls (donor only), FIT-GFP or FITmSS271AA-GFP (donor) and bHLH039-mCherry (acceptor) as FRET pair. GFP was excited with a linearly polarized pulsed (32 MHz) 485 nm laser and 0.01-0.1 µW output power. Measurements were recorded with a UPLSAPO 60XW (Olympus) objective, zoom factor of 8, pixel dwell time of 12.5 µs, objective frame size of 256x256, and 60 frames. Measurements were controlled with the manufacturer’s FV31S-SW (Olympus) software and SymPhoTime 64 (PicoQuant) software. SymPhoTime 64 (PicoQuant) software was used for analysis in the respective regions of interest (whole nucleus, NB, NP) and to generate color-coded fluorescence lifetime value images. Number of parameters for the fit depended on the region of interest.

### Circularity quantification

Circularity quantification was performed with the software ImageJ (National Institutes of Health). Full intensity projection images were generated from Z-stacks in the ZEN lite (Zeiss) software and exported as TIFF (no compression, all dimensions). Images were duplicated in ImageJ and converted to RGB and 8-bit. Correct scale was set (in µm) under ‘Analyze’ - ‘Set Scale’. Threshold for the intensity limit (areas below that limit were not considered for quantification) was set under ‘Image’ - ‘Adjust’ - ‘Threshold’ and was set manually for every image. To separate the nuclear bodies better, ‘Process’ - ‘Binary’ - ‘Watershed’ was used. Parameters that should be quantified were selected under ‘Analyze’ - ‘Set Measurements’. To perform the analysis, ‘Analyze’ - ‘Analyze Particles’ was selected. Calculated values were further processed in Excel (Microsoft Corporation).

### Nuclear body quantification

Nuclear body quantification was performed with the software ImageJ (National Institutes of Health) and additional plugin ‘3D Object Counter’. Z-stacks were exported from the ZEN lite (Zeiss) software as TIFF (no compression, all dimensions) first. In ImageJ, Z-stacks were converted to RGB and 8-bit. Correct scale was set (in µm) under ‘Properties’. Parameters that should be quantified were selected under ‘Plugins’ - ‘3D Object Counter’ - ‘Set 3D Measurements’. To perform the analysis, ‘Plugins’ - ‘3D Object Counter’ - ‘3D object counter’ was selected. Threshold for the intensity limit (areas below that limit were not considered for quantification) was set manually for every z-stack. Calculated values were further processed in Excel (Microsoft Corporation). Only size between 0,01-15 µm³ was considered.

### Protein domain prediction

IDRs in FIT/FITmSS271AA were predicted with the tool PONDR-VLXT (www.pondr.com, Molecular Kinetics, Inc.). According to the sequence of the protein, a PONDR score was determined for each amino acid. A score above 0.5 indicates intrinsic disorder. The bHLH domain of FIT was predicted with InterPro (www.ebi.ac.uk/interpro, EMBL-EBI).

### Statistical analysis

Line and bar diagrams represent the mean and standard deviation. Box plots show 25-75 percentile with min-max whiskers, mean as small square and median as line. Graphs and statistical analysis were created and performed with OriginPro (OriginLab Corporation). Data was tested for normal distribution with the Shapiro-Wilk test. Statistical significance of data with normal distribution was tested by one-way Anova with Tukey post-hoc test. Statistical significance of data with non-normal distribution was tested by Mann-Whitney test. Different letters indicate statistically significant differences (P < 0.05). Illustrations were created with BioRender.com.

### Accession numbers

Sequence data from this article can be found in the EMBL/GenBank data libraries under accession numbers*: bHLH039* (AT3G56980), *COILIN* (AT1G13030), *FIT* (AT2G28160), *FRO2* (AT1G01580), *IRT1* (AT4G19690), *P15H1* (AT1G11570), *PIF3* (AT1G09530), *PIF4* (AT2G43010), *PININ* (AT1G15200), *SR45* (AT1G16610), *SRm102* (AT2G29210), *U2B”* (AT2G30260), *UAP56H2* (AT5G11170), and *ZAT12* (AT5G59820).

## Supplemental Data

**Supplemental Figure S1.** FIT NBs induced by blue light and a standardized FIT NB analysis procedure was developed to analyze the characteristics and dynamics of FIT NBs. (Supports Figure 1)

**Supplemental Figure S2.** An intrinsically disordered region, IDR^Ser271/272^, is present in the FIT C-terminus and disrupted in the FITmSS271AA mutant. (Supports Figure 2, 3, and 4)

**Supplemental Figure S3.** FIT NBs did not colocalize with Cajal body components (designated type I). (Supports Figure 5 and 6)

**Supplemental Figure S4.** Type II and III NB markers are similarly localized upon single expression as upon co-expression with FIT, except PININ. (Supports Figure 5 and 6)

**Supplemental Figure S5.** PB markers are similarly localized upon single expression and upon co-expression with FIT. (Supports Figure 7)

**Supplemental Figure S6.** Differential expression of intron retention splicing variants of iron deficiency genes in response to iron deficiency and blue light. (Supports Figure 9)

**Supplemental Figure S7.** The variations in gene expression levels between white light and blue light are consistent across both intron retention splicing variants and the overall transcript pool. (Supports Figure 9 and Supplemental Figure S6)

**Supplemental Table S1.** List of primers used in this study.

**Supplemental Table S2.** List of vectors used in this study.

**Supplemental Movie S1.** Light induction triggers the formation of NBs with FIT and FITmSS271AA with different dynamics. (Supports Figure 1 and 2)

## Abbreviations

bHLH: basic helix-loop-helix
bHLH039: BASIC HELIX-LOOP-HELIX039
C: mCherry
FIT: FER-LIKE IRON DEFICIENCY-INDUCED TRANSCRIPTION FACTOR
FLIM: fluorescence lifetime imaging microscopy
FRAP: fluorescence recovery after photobleaching
FRET: fluorescence resonance energy transfer
G: GFP
GFP: GREEN FLUORESCENT PROTEIN
IDR: intrinsically disordered region
IR: intron retention
LLPS: liquid-liquid phase separation
mCherry: monomeric Cherry
mRFP: monomeric RED FLUORESCENT PROTEIN
NB: nuclear body
NP: nucleoplasm
PB: photobody
R: mRFP
TF: transcription factor

## Acknowledgements

We thank Elke Wieneke and Monique Eutebach (Institute of Botany, Heinrich-Heine-University, Düsseldorf, Germany) for excellent technical assistance. We acknowledge the contribution of the student Daniel Torno. We thank Rebecca C. Burkart (Institute of Developmental Genetics, Heinrich-Heine-University, Düsseldorf, Germany), Sebastian Hänsch and Stefanie Weidtkamp-Peters (Center for Advanced Imaging, Heinrich-Heine-University, Düsseldorf, Germany) for help and advice with microscopy and data analysis. K.T. is an associate member of the DFG-funded International Graduate School for Plant Science iGRAD*plant* (IRTG 2466 “Network, exchange, and training program to understand plant resource allocation”), Düsseldorf, Germany.

