## Supplemental Movie S1 for "FER-LIKE IRON DEFICIENCY-INDUCED TRANSCRIPTION FACTOR (FIT) accumulates in homo- and heterodimeric complexes in dynamic and inducible nuclear condensates associated with speckle components"

### Slide 1
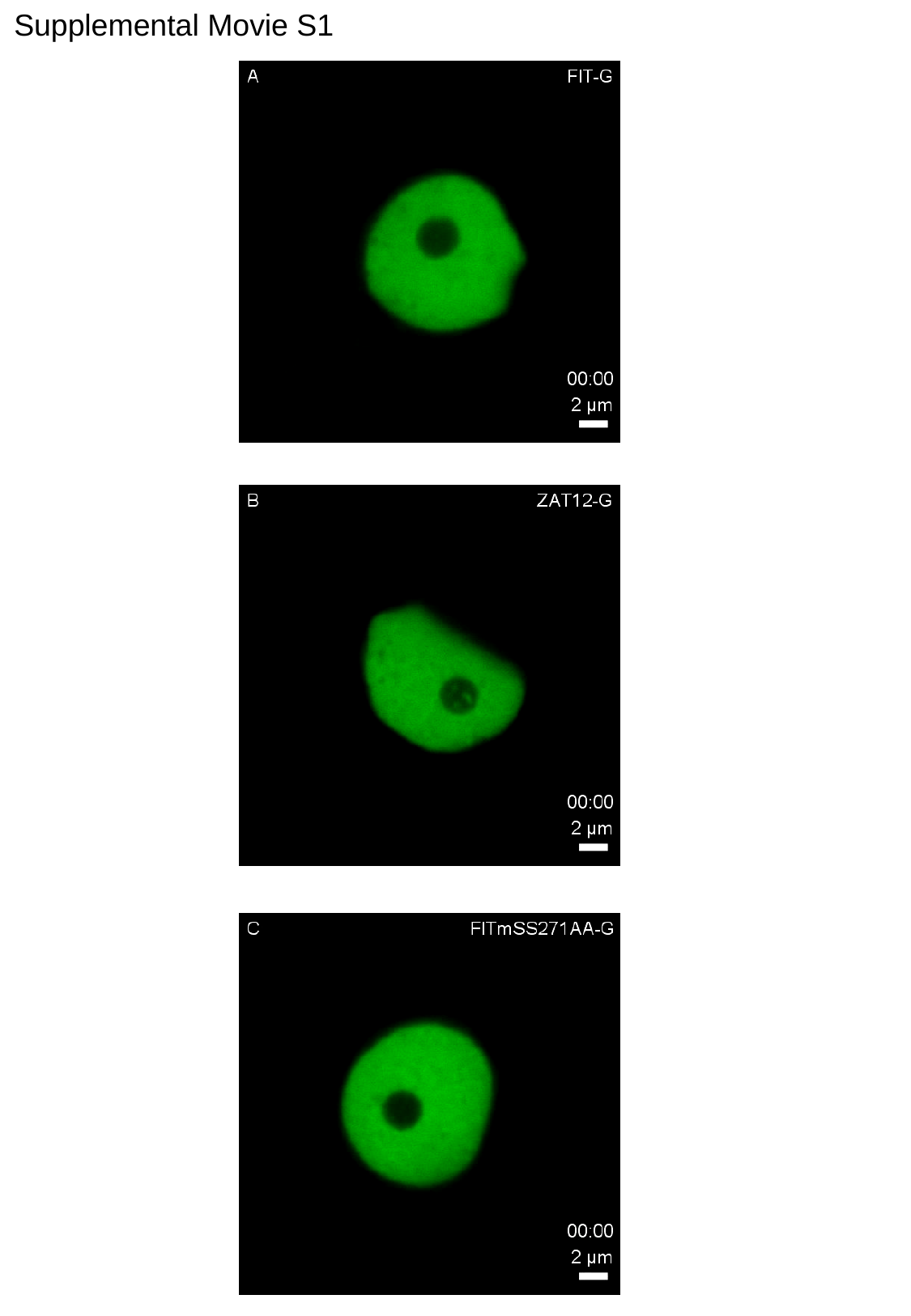

Supplemental Movie S1

### Slide 2
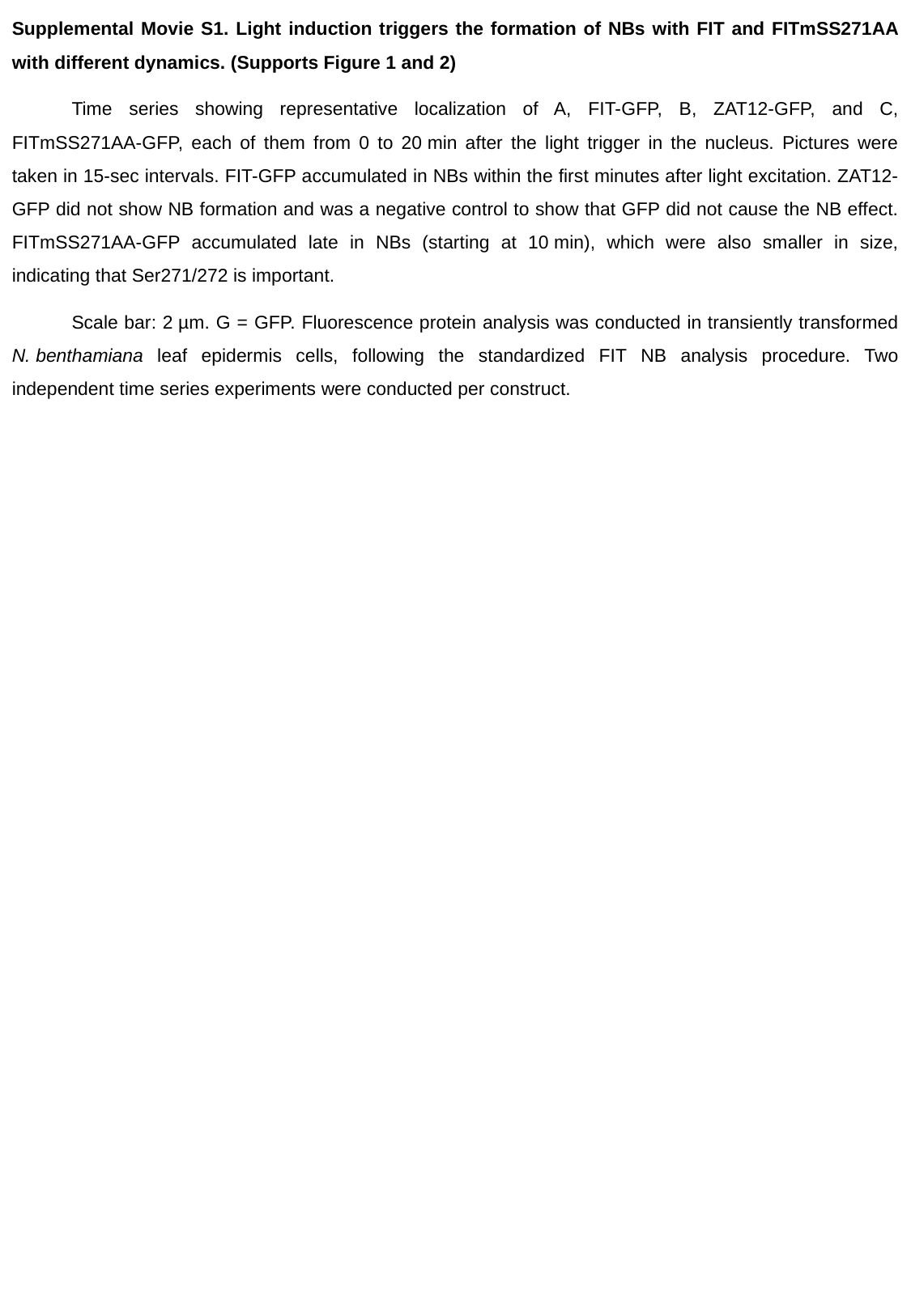

Supplemental Movie S1. Light induction triggers the formation of NBs with FIT and FITmSS271AA with different dynamics. (Supports Figure 1 and 2)
Time series showing representative localization of A, FIT-GFP, B, ZAT12-GFP, and C, FITmSS271AA-GFP, each of them from 0 to 20 min after the light trigger in the nucleus. Pictures were taken in 15-sec intervals. FIT-GFP accumulated in NBs within the first minutes after light excitation. ZAT12-GFP did not show NB formation and was a negative control to show that GFP did not cause the NB effect. FITmSS271AA-GFP accumulated late in NBs (starting at 10 min), which were also smaller in size, indicating that Ser271/272 is important.
Scale bar: 2 µm. G = GFP. Fluorescence protein analysis was conducted in transiently transformed N. benthamiana leaf epidermis cells, following the standardized FIT NB analysis procedure. Two independent time series experiments were conducted per construct.
