## Supplemental Table S1 for "FER-LIKE IRON DEFICIENCY-INDUCED TRANSCRIPTION FACTOR (FIT) accumulates in homo- and heterodimeric complexes in dynamic and inducible nuclear condensates associated with speckle components"

**Supplemental Table S1. List of primers used in this study.**
All primers originated in this study

| Primer name | Sequence 5' → 3' | Purpose |
|---|---|---|
| pFIT GG fw | AAAGGTCTCAACCTCTAAAGATGTGCTGATAAGT | GreenGate Cloning *FIT* promoter |
| pFIT GG rv | AAAGGTCTCATGTTTTGTGTGTTTTTGTC | GreenGate Cloning *FIT* promoter |
| cFIT GG fw | AAAGGTCTCAGGCTTAATGGAAGGAAGAGT | GreenGate Cloning *FIT* CDS |
| cFIT GG rv | AAAGGTCTCACTGAAGTAAATGACTTGATG | GreenGate Cloning *FIT* CDS |
| PIF3 GW fw | GGGGACAAGTTTGTACAAAAAAGCAGGCTATGC CTCTGTTTGAGCTT | Gateway Cloning *PIF3* CDS |
| PIF3 GW rv | GGGGACCACTTTGTACAAGAAAGCTGGGTCCGA CGATCCACAAAACTG | Gateway Cloning *PIF3* CDS |
| PIF4 GW fw | GGGGACAAGTTTGTACAAAAAAGCAGGCTATGG AACACCAAGGTTGG | Gateway Cloning *PIF4* CDS |
| PIF4 GW rv | GGGGACCACTTTGTACAAGAAAGCTGGGTCGTG GTCCAAACGAGAACC | Gateway Cloning *PIF4* CDS |
| cEF fw | TATGGGATCAAGAAACTCACAAT | *EF1Balpha* RT-qPCR |
| cEF rv | CTGGATGTACTCGTTGTTAGGC | *EF1Balpha* RT-qPCR |
| gEF fw | TCCGAACAATACCAGAACTAC | *EF1Balpha* (genomic) RT-qPCR |
| gEF rv | CCGGGACATATGGAGGTAAG | *EF1Balpha* (genomic) RT-qPCR |
| FIT fw | CCCTGTTTCATAGACGAGAACC | *FIT* RT-qPCR |
| FIT rv | ATCCTTCATACGCCCTCTCC | *FIT* RT-qPCR |
| FIT IR1 fw | CAACGACGGTGATGATTCTTC | *FIT intron 1* RT-qPCR |
| FIT IR1 rv | CAATATTTACAGTTTCATTAGTCATTACCA | *FIT intron 1* RT-qPCR |
| FIT IR2 fw | ACCTAAAAAAACCAAATCAATCTTC | *FIT intron 2* RT-qPCR |
| FIT IR2 rv | GGAGAAGGAGAGCTTAGGTTAGA | *FIT intron 2* RT-qPCR |
| bHLH39 fw | GACGGTTTCTCGAAGCTTG | *BHLH039* RT-qPCR |
| bHLH039 rv | GGTGGCTGCTTAACGTAACAT | *BHLH039* RT-qPCR |
| bHLH39 IR1 fw | GATTAGACCCGTTATAGTTTATGCAT | *BHLH039 intron 1* RT-qPCR |
| bHLH039 IR1 rv | TATTAGCTTCTTCACTTGCTCTTGC | *BHLH039 intron 1* RT-qPCR |
| IRT1 fw | AAGCTTTGATCACGGTTGG | *IRT1* RT-qPCR |
| IRT1 rv | TTAGGTCCCATGAACTCCG | *IRT1* RT-qPCR |
| IRT1 IR2 fw | TGGGATCATAGTTCACTCGGTGG | *IRT1 intron 2* RT-qPCR |
| IRT1 IR2 rv | TATATACAATTTCAACATTTGGCTAAAGAT | *IRT1 intron 2* RT-qPCR |
| FRO2 fw | TTAGGTCCCATGAACTCCG | *FRO2* RT-qPCR |
| FRO2 rv | AAGATGTTGGAGATGGACGG | *FRO2* RT-qPCR |
| FRO2 IR3 fw | GTAGATTCTGTGCTTGATTTCTTTTTAAAA | *FRO2 intron 3* RT-qPCR |
| FRO2 IR3 rv | GGTTAATCCCATTGCCGGTAG | *FRO2 intron 3* RT-qPCR |
| FRO2 IR4 fw | CTACCGGCAATGGGATTAACC | *FRO2 intron 4* RT-qPCR |
| FRO2 IR4 rv | GCTCCATTCAGAGTGACTACTTTA | *FRO2 intron 4* RT-qPCR |
| FRO2 IR5 fw | TATTGATCTAATTCTAAACTATTCTTTTTG | *FRO2 intron 5* RT-qPCR |
| FRO2 IR5 rv | CTACAGAAACAGCAAGTCGGT | *FRO2 intron 5* RT-qPCR |
