## Supplemental Table S2 for "FER-LIKE IRON DEFICIENCY-INDUCED TRANSCRIPTION FACTOR (FIT) accumulates in homo- and heterodimeric complexes in dynamic and inducible nuclear condensates associated with speckle components"

**Supplemental Table S2. List of vectors used in this study.**

| Vector | Application | Source |
|---|---|---|
| ipABind:cFIT-GFP | Imaging, FRAP, anisotropy, FRET-FLIM | Gratz et al., 2019 |
| ipABind:cFIT-mCherry | Imaging | |
| ipABind:cFITmSS271AA-GFP | Imaging, anisotropy, FRET-FLIM | |
| ipABind:cbHLH039-mCherry | Imaging, FRET-FLIM | Trofimov et al., 2019 |
| pMDC83:ZAT12-GFP | Imaging | Le et al., 2016 |
| pROK2:COILIN-mRFP | Imaging | The Plant Nuclear Marker collection (NASC) |
| pROK2:P15H1-mRFP | Imaging | |
| pROK2:PININ-mRFP | Imaging | |
| pROK2:SR45-mRFP | Imaging | |
| pROK2:SRm102-mRFP | Imaging | |
| pROK2:U2B“-mRFP | Imaging | |
| pROK2:UAP56H2-mRFP | Imaging | |
| ipABind:cPIF3-mCherry | Imaging | this study |
| ipABind:cPIF4-mCherry | Imaging | |
