## Supplemental Figure S1 for "FER-LIKE IRON DEFICIENCY-INDUCED TRANSCRIPTION FACTOR (FIT) accumulates in homo- and heterodimeric complexes in dynamic and inducible nuclear condensates associated with speckle components"

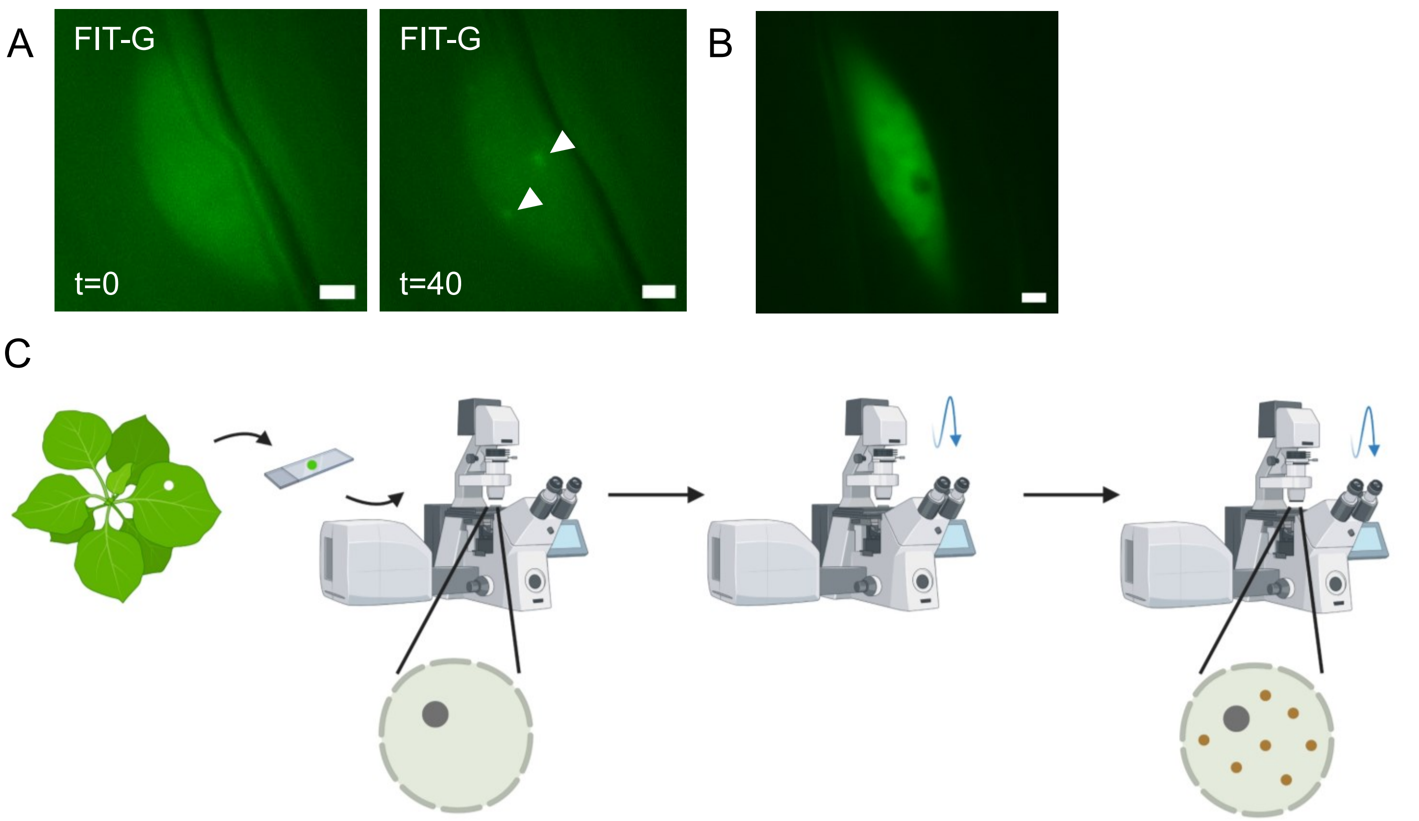


Induction of protein expression in *Nicotiana benthamiana* 16 h before imaging.

Data acquisition t=0. Check expression of construct, take images and measurements before NB formation.

Light induction of NB formation by excitation with 488 nm laser for 1 min.

Data acquisition t=5. Take images and measurements after NB formation.

**Supplemental Figure S1. FIT NBs induced by blue light and a standardized FIT NB analysis procedure was developed to analyze the characteristics and dynamics of FIT NBs. (Supports Figure 1)**

A, Induction of FIT NBs in Arabidopsis root epidermis cells of the root differentiation zone at t=0 and t=40 min of 5-d-old seedling (2x35$S_{pro}$:FIT-GFP) grown under iron deficiency. FIT-GFP signals were evenly distributed in the nucleus at t=0 min, and after induction by excitation with 488 nm laser NB formation accumulated in NBs at t=40 min. Root epidermis cells developed few NBs with weak FIT-GFP signals, sometimes taking up to two hours to appear. Representative pictures from four independent experiments. B, Arabidopsis root epidermal cells of the root differentiation zone of 5-d-old seedling (proFIT:FIT-GFP) grown under iron deficiency do not show NBs when taken directly from white light. Representative pictures from three independent experiments. Scale bar: 2 μm. C, Experimental steps for FIT NB induction in transiently transformed *N. benthamiana* leaf epidermis cells. Fluorescence protein expression was induced by β-estradiol ('induction of protein expression') 16 h prior to imaging and measurements. Leaf discs were excised, and initial fluorescence images and measurements were taken ('data acquisition t=0'). Leaf discs were exposed to 488 nm laser light as a light trigger for 1 min ('light induction of NB formation'), and 5 min later, fluorescence images and measurements were taken again ('data acquisition t=5'). With this procedure, FIT NBs were visible, and their characteristics could be analyzed. In some cases, fluorescence images and measurements were taken at t=15 min, as indicated in the text. Imaging was performed at the respective wavelengths for detection of GFP and mRFP/mCherry, respectively.
