## Supplemental Figure S2 for "FER-LIKE IRON DEFICIENCY-INDUCED TRANSCRIPTION FACTOR (FIT) accumulates in homo- and heterodimeric complexes in dynamic and inducible nuclear condensates associated with speckle components"

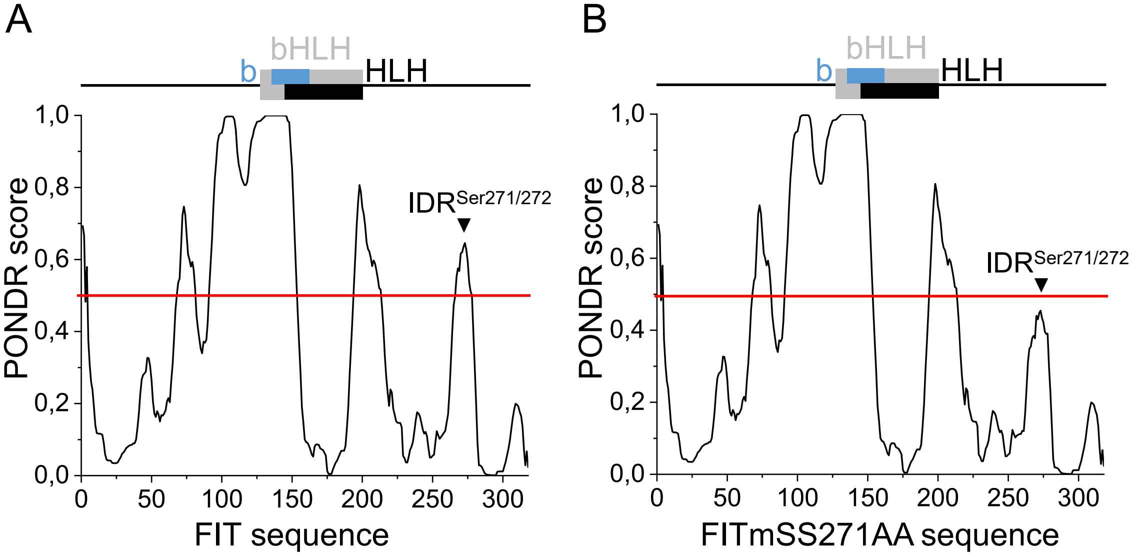

**Supplemental Figure S2. An intrinsically disordered region, IDR$^{Ser271/272}$, is present in the FIT C-terminus and disrupted in the FITmSS271AA mutant. (Supports Figure 2, 3, and 4)**

Diagrams representing the PONDR scores for each amino acid position in A, FIT, and B, FITmSS271AA protein sequences. Analysis was performed via the tool PONDR-VLXT (Molecular Kinetics, Inc.). A score >0.5 indicates intrinsic disorder. The 0.5 threshold is marked with a red line. Above the graph, schematic representation of FIT protein showing the position of the bHLH domain in grey (126-201 aa) and subdivided into the basic region in blue (DNA binding site, 132-162 aa) and the helix-loop-helix region in black (dimerization site, 142-201 aa). Domain prediction was performed with InterPro (EMBL-EBI). FIT has four regions with a score >0.5 that are predicted IDRs, two of them in the C-terminal part following the bHLH domain, with one out of them comprising the position SS271/272, indicated by an arrowhead, termed IDR$^{Ser271/272}$. In FITmSS271AA, the PONDR score dropped for this region below the threshold.
