## Supplemental Figure S3 for "FER-LIKE IRON DEFICIENCY-INDUCED TRANSCRIPTION FACTOR (FIT) accumulates in homo- and heterodimeric complexes in dynamic and inducible nuclear condensates associated with speckle components"

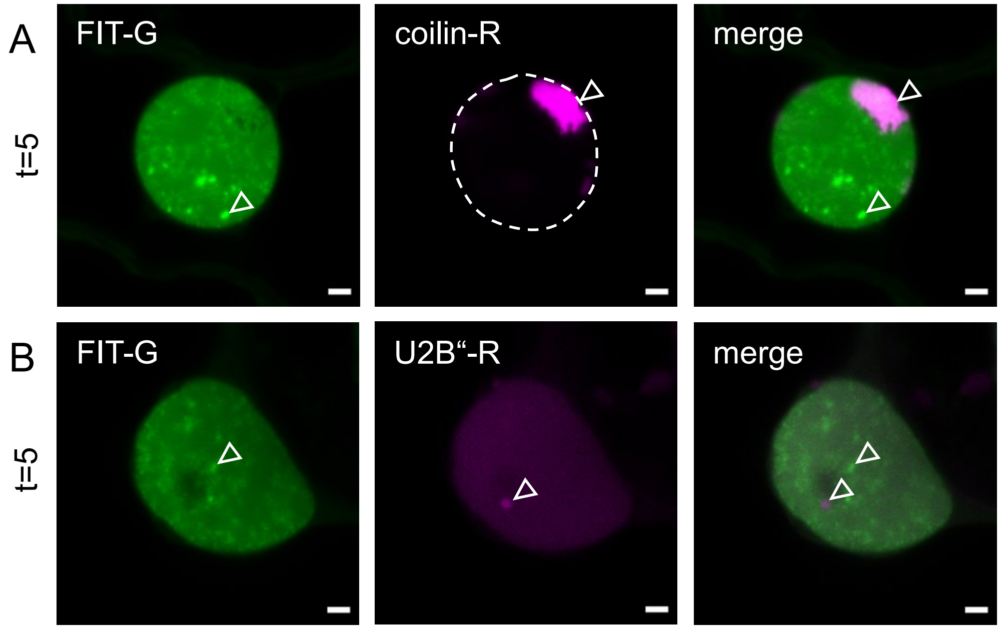

**Supplemental Figure S3. FIT NBs did not colocalize with Cajal body components (designated type I). (Supports Figure 5 and 6)**

Confocal images showing localization of FIT-GFP and NB markers (type I) upon co-expression in the nucleus at t=5 min. Co-expression of FIT-GFP with A, coilin-mRFP, and B, U2B”-mRFP. FIT NBs were present at t=5 min and did not colocalize with NBs of the two markers.

Scale bar: 2 µm. Arrowheads indicate non colocalizing NBs. G = GFP; R = mRFP. Fluorescence protein analysis was conducted in transiently transformed *N. benthamiana* leaf epidermis cells, following the standardized FIT NB analysis procedure. In all examined cells, the proteins showed no colocalization. Representative images from three independent experiments.
