## Supplemental Figure S4 for "FER-LIKE IRON DEFICIENCY-INDUCED TRANSCRIPTION FACTOR (FIT) accumulates in homo- and heterodimeric complexes in dynamic and inducible nuclear condensates associated with speckle components"

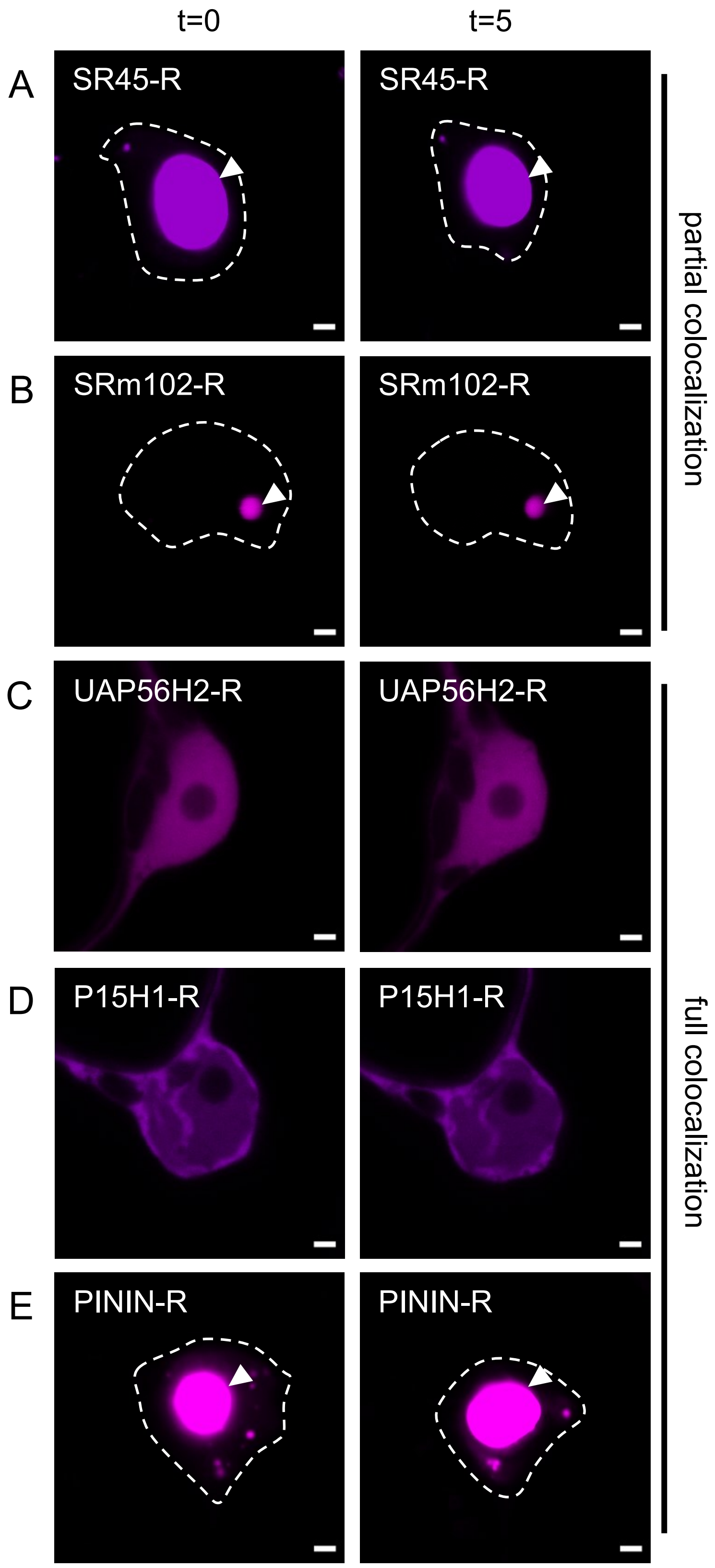

**Supplemental Figure S4. Type II and III NB markers are similarly localized upon single expression as upon co-expression with FIT, except PININ. (Supports Figure 5 and 6)**

Confocal images showing localization of NB markers (type II and III) upon their single expression in the nucleus at t=0 and t=5 min, in A, SR45-mRFP, B, SRm102-RFP, C, UAP56H2-mRFP, D, P15H1-mRFP, and E, PININ-mRFP. Single SR45-mRFP and SRm102-RFP localized in NBs similar to the colocalization with FIT at t=0 and t=5 min (compare with **Figure 5**). Single UAP56H2-mRFP and P15H1-mRFP did not localize in NBs and were uniformly distributed, similar to the colocalization with FIT at t=0 (compare with **Figure 6, A and B**). Only single PININ-mRFP showed a different localization pattern between its single expression versus co-expression with FIT-GFP. Upon single expression it localized in NBs at t=0 and t=5 min, while in co-expression with FIT-GFP it showed no NBs at t=0 but followed the FIT NB pattern at t=5 min (compare with **Figure 6C**).

Scale bar: 2 µm. Arrowheads indicate NBs. R = mRFP. Fluorescence protein analysis was conducted in transiently transformed *N. benthamiana* leaf epidermis cells, following the standardized FIT NB analysis procedure. Representative images from three to five independent experiments.
