## Supplemental Figure S5 for "FER-LIKE IRON DEFICIENCY-INDUCED TRANSCRIPTION FACTOR (FIT) accumulates in homo- and heterodimeric complexes in dynamic and inducible nuclear condensates associated with speckle components"

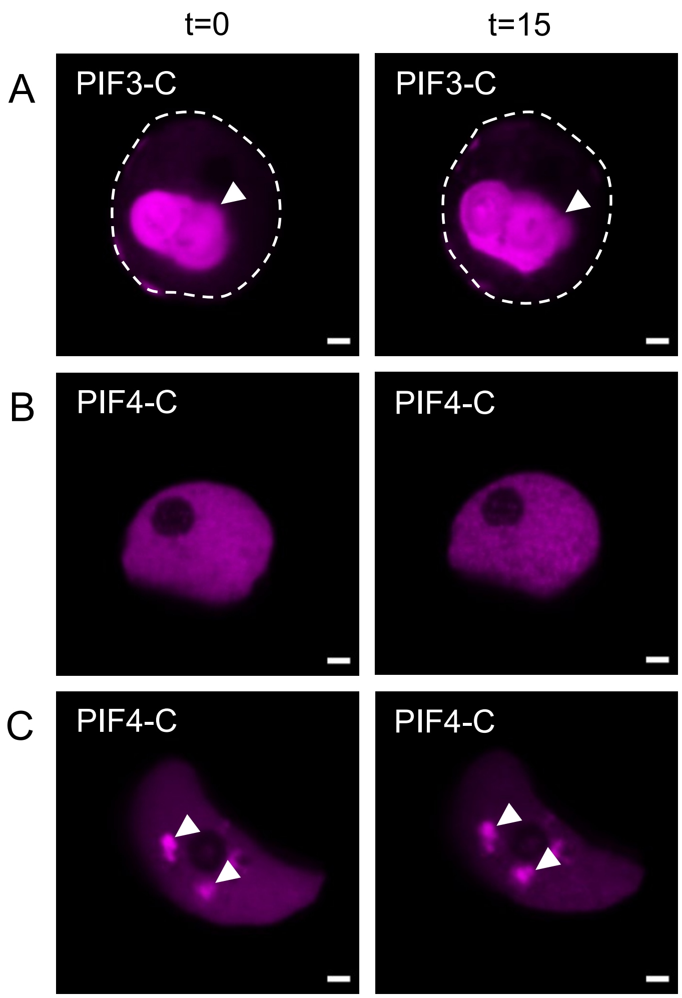

**Supplemental Figure S5. PB markers are similarly localized upon single expression and upon co-expression with FIT. (Supports Figure 7)**

Confocal images showing localization of PB markers upon their single expression in the nucleus at t=0 and t=15 min, in A, PIF3-mCherry, and in B and C, PIF4-mCherry in two different patterns. Single PIF3-mCherry localized to a very large PB at t=0 and t=15 min. Single PIF4-mCherry localized either in a uniform manner in the nucleus as seen in B, or in several PBs as seen in C. Hence, PIF3-mCherry and PIF4-mCherry were similarly localized in single expression as upon co-expression with FIT-GFP (compare with **Figure 7**).

Scale bar: 2 µm. Arrowheads indicate NBs. C = mCherry. Fluorescence protein analysis was conducted in transiently transformed *N. benthamiana* leaf epidermis cells, following the standardized FIT NB analysis procedure. Representative images from three to six independent experiments.
