## Supplemental Figure S6 for "FER-LIKE IRON DEFICIENCY-INDUCED TRANSCRIPTION FACTOR (FIT) accumulates in homo- and heterodimeric complexes in dynamic and inducible nuclear condensates associated with speckle components"

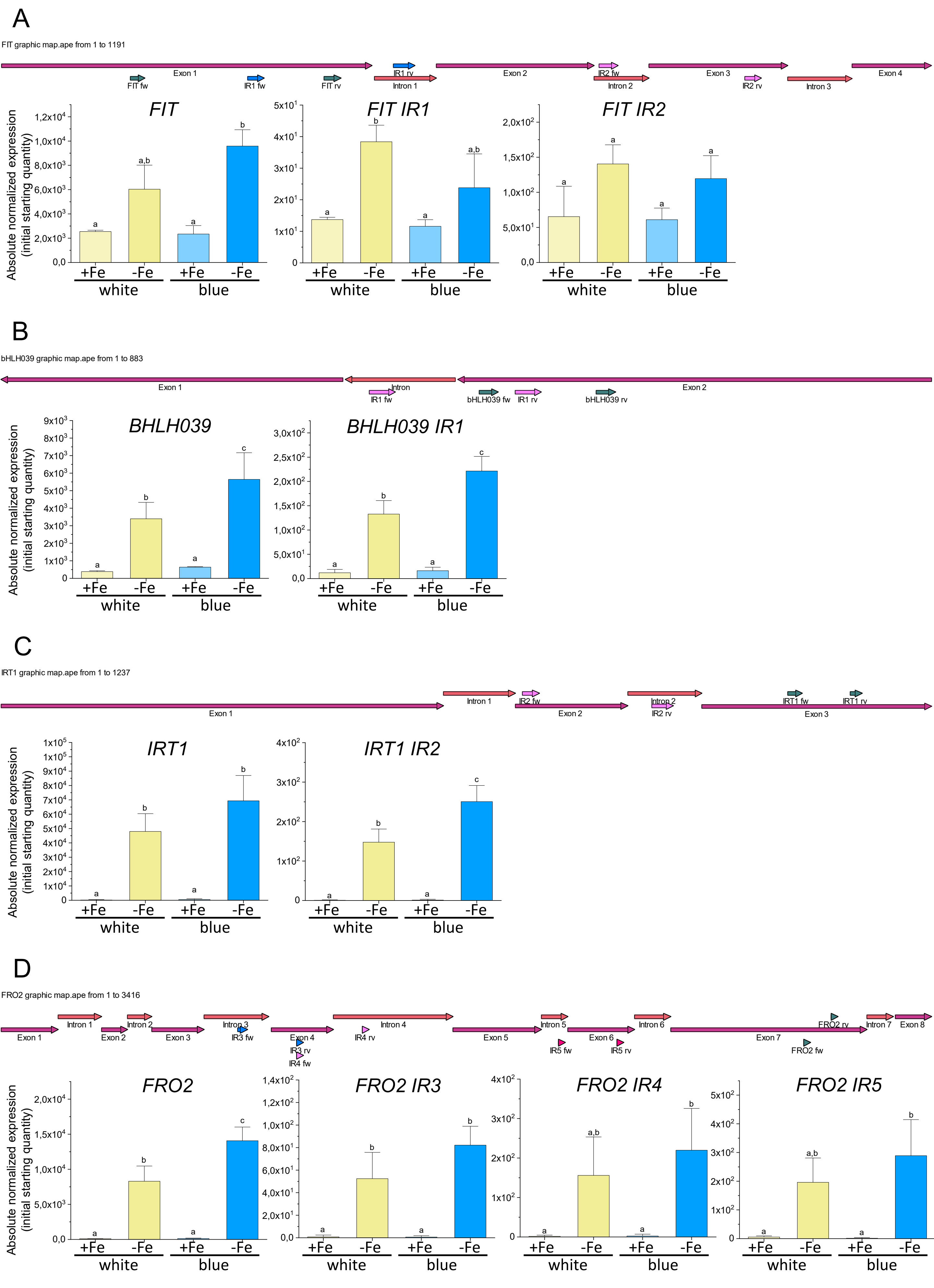

**Supplemental Figure S6. Differential expression of intron retention splicing variants of iron deficiency genes in response to iron deficiency and blue light. (Supports Figure 9)**

Gene expression analysis of total transcript abundance of iron deficiency genes A, *FIT*, B, *BHLH039*, C, *IRT1*, and D, *FRO2* and selected transcripts with intron retention (IR) splicing variants previously reported for these genes (Li et al., 2013). 5-d-old Arabidopsis seedlings grown under white light for 5 d under iron deficient and iron sufficient conditions were exposed for 1.5-2 h to blue light and in parallel as control to white light. At the top of A-D, overview of exon-intron structures and sites of qPCR primers, detecting IR variant transcripts and respective total amounts of transcripts. Bottom, gene expression data for the indicated gene products. The absolute expression levels of IR variant transcripts were at least 20-40 times lower than those of total transcripts. A, Due to the low abundance of FIT IR splicing variants, there was only one case where gene expression was significantly changed in response to an environmental treatment. *FIT IR1* was upregulated under iron deficiency versus sufficiency in blue light, similar to *FIT*. There was no significant difference for *FIT IR2* splicing variant. B, *BHLH039* and *BHLH039 IR1* splicing variant gene expression increased significantly in response to low iron supply and after blue light exposure compared to the white light control. C, *IRT1* and *IRT1 IR2* splicing variant gene expression was significantly higher under iron deficient versus sufficient conditions. Gene induction in response to low iron supply did not change after blue light exposure compared to the white light control but was increased for *IRT1 IR2* splicing variant. D, *FRO2* gene expression was enhanced under blue light versus white light in iron deficient conditions. The three *FRO2 IR3-5* splicing variants were more abundant under iron deficiency than sufficiency, but not differently regulated between white and blue light.

Four experiments were conducted, one representative result is shown. Bar diagrams represent the mean and standard deviation of three replicates with twenty seedlings and two technical replicates each (n=3). Statistical analysis was performed with one-way ANOVA and Tukey post-hoc test. Different letters indicate statistically significant differences ($P < 0.05$).
