## Supplemental Figure S7 for "FER-LIKE IRON DEFICIENCY-INDUCED TRANSCRIPTION FACTOR (FIT) accumulates in homo- and heterodimeric complexes in dynamic and inducible nuclear condensates associated with speckle components"

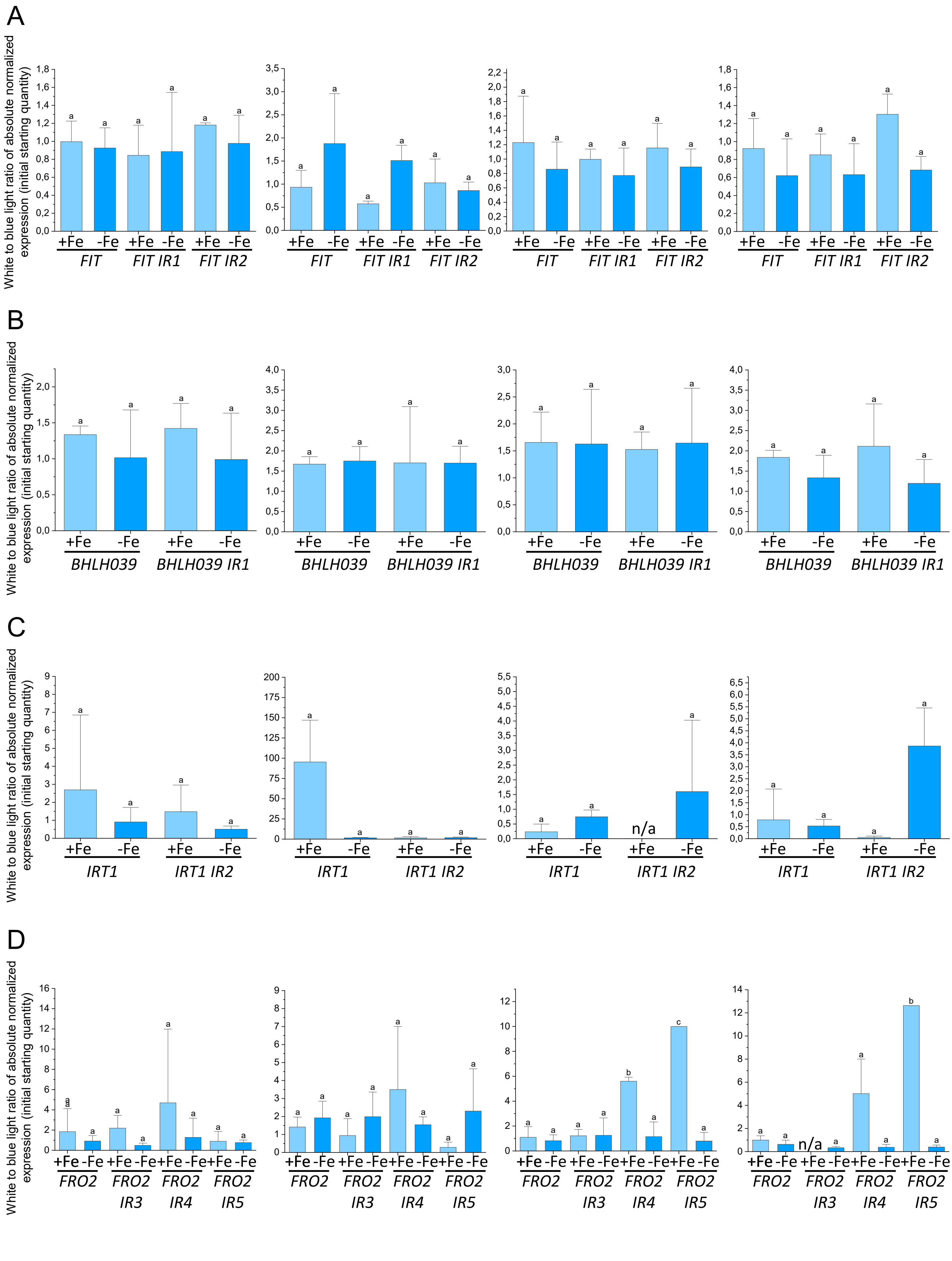

**Supplemental Figure S7. The variations in gene expression levels between white light and blue light are consistent across both intron retention splicing variants and the overall transcript pool. (Supports Figure 9 and Supplemental Figure S6)**

Ratios of absolute normalized gene expression levels in white light versus blue light for total transcript abundance of iron deficiency genes A, *FIT*, B, *BHLH039*, C, *IRT1*, and D, *FRO2* and of previously reported intron retention (IR) splicing variants (Li et al., 2013). Respective absolute normalized gene expression levels and explanations about the experiment are represented in **Supplemental Figure S6**. The ratios obtained for the four conducted experiments are presented from left to right. No differences in the ratios were found between the transcript abundance of IR spicing variants versus total transcript abundance.

Bar diagrams represent the mean and standard deviation of three ratios (n=3). Statistical analysis was performed with one-way ANOVA and Tukey post-hoc test. Different letters indicate statistically significant differences ($P < 0.05$). n/a = no gene expression value.
